# Evaluating Limits of Machine Learning-Assisted Raman Spectroscopy in Classification of Biological Samples

**DOI:** 10.64898/2026.02.26.708284

**Authors:** Aman Yadav, Arlin Birkby, Noah Armstrong, Assame Arnob, Ming-Hsun Chou, Alma Fernandez, Aart J. Verhoef, Zhenhuan Yi, Siddhant Gulati, Siddhi Kotnis, Qing Sun, Katy C. Kao, Hung-Jen Wu

## Abstract

Machine learning (ML)-assisted Raman spectroscopy has become a powerful analytical tool for the classification and identification of analytes; however, technical challenges impacting its detection accuracy have not been investigated. This study explores experimental factors affecting classification performance. Among the evaluated ML models, ML algorithms show minimal impacts on classification accuracy. Instead, experimental factors, including spectral similarity between tested samples and the data quality, dominate detection performance. Increases in spectral noises and spectral similarity significantly reduce classification accuracy. In well-controlled samples with low experimental noise, ML-assisted Raman spectroscopy can discriminate lipid mixtures with a composition difference of 1.85 mol%. To assess the effect of biological heterogeneity, we analyzed single-cell Raman spectra from *Saccharomyces cerevisiae* strains carrying single, double, or triple gene mutations. Intrinsic cell-to-cell variability introduced substantial spectral differences, severely reducing the accuracy of multiclass classification of these genetically similar strains at the single-cell level. Averaging Raman spectra across multiple cells improved classification accuracy by reducing this spectral variability. We also assess the effectiveness of transfer learning across different Raman spectrometers, specifically by applying a ML model trained on one instrument to another Raman spectrometer. Transfer learning can be improved with proper instrument calibration, highlighting the importance of instrument standardization. Overall, our results demonstrate that data quality and spectral similarity are the primary bottlenecks in ML-assisted Raman spectroscopy. Careful attention to sample preparation, data acquisition, measurement conditions, and instrument calibration is critical to achieving robust and reliable classification performance.

## Introduction

Raman scattering is an inelastic light scattering phenomenon that could be used to detect molecular vibrations. Due to unique molecular characteristics, Raman spectroscopy provides a fingerprint identification of the analytical samples ^1–3^. Raman spectroscopy is a fast, non-destructive, and real-time measurement technique widely utilized across various fields. However, its application is often constrained by the complexity of spectral data. In recent years, machine learning (ML)-assisted Raman spectroscopy has become a robust analytical tool, benefiting from advancements in machine learning techniques.

Subtle variations in Raman spectra are often challenging to detect. The experimental factors like instrumental variations, measurement errors, or artifacts caused by fluorescence background can influence the spectral analysis ^4^. Machine learning techniques have been used to extract meaningful information from spectral data more effectively ^5^. Multiple supervised (Partial Least Squares (PLS), Linear Discriminant Analysis (LDA)) and unsupervised (Principal Component Analysis (PCA), Autoencoder(AE)) algorithms have been widely used for dimensionality reduction ^6^. PCA is commonly used to reduce data collinearity and the number of variables, which allows algorithms to deliver results quickly and efficiently when analyzing similar spectra within a class. LDA is based on extracting different hyperplanes, or linear functions, that effectively enable discrimination between two or more classes in multivariate space ^7^. These methods have been widely used for dimensionality reduction in Raman spectral data. ML algorithms, such as Support Vector Machine (SVM), Random Forest (RF), K-Nearest Neighbors (KNN), Decision Tree (DT), and Neural Networks (NN), have demonstrated improved performance when combined with Principal Component Analysis (PCA) and Linear Discriminant Analysis (LDA). Further, the use of ML algorithms like Convolutional Neural Networks (CNNs) allows them to autonomously learn features by optimizing network weights, which helps to remove the use of methods that require human-defined feature definitions ^5^. ML-assisted Raman spectroscopy has broad applications in forensics, pharmaceuticals, food industry, materials science, and biomedical research ^8–16^.

The effectiveness of sample classification using ML-assisted Raman spectroscopy depends on several factors: the choice of ML algorithms, spectral similarity, and data quality. Data quality could be shaped by experimental conditions, including sample preparation, natural variations in biological samples, and inconsistencies arising from different instruments, laboratories, or technicians. ^4, 17^. Spectral variations, caused by environmental factors (e.g., room lighting, background signals, fluorescence) and instrumental noise (e.g., dark current, photon shot, read noise) ^18, 19^, can reduce the signal-to-noise ratio (SNR) and hinder classification accuracy ^20^. Moreover, high spectral similarity between samples can further challenge ML-based Raman spectroscopy classification.

This study investigated factors affecting classification accuracy in ML assisted Raman spectroscopy. We evaluated the performance of ML algorithms widely used in the literature, including Naïve Bayes (Gaussian), Support Vector Machine (SVM), K-Nearest Neighbors (KNN), Neural Networks (NN), and Convolutional Neural Networks (CNN). We prepared samples with varying spectral similarity by titrating octanoic acid (OA), a medium-chain (C8) saturated fatty acid, into glyceryl trioctanoate (GTO), a triglyceride derived from OA. Due to their similar chemical structures, GTO and OA exhibit comparable Raman spectra, enabling us to assess the effect of spectral similarity on classification accuracy. We also evaluated the impact of intra-day and inter-day variations on ML performance. To further explore the relationships among spectral variations, similarity, and accuracy, simulated noise was introduced into the spectral data. Additionally, we developed an instrument calibration technique to facilitate transfer learning across different Raman spectrometers. The influence of cell-to-cell variations was examined using spectral data from *Saccharomyces cerevisiae* with single, double, and triple gene mutations. Our findings indicate that ML algorithms had minimal impact on classification accuracy, whereas data quality significantly influenced the performance of ML-assisted Raman spectroscopy. Careful planning of sample collection, data acquisition, and instrument calibration is essential for enhancing Raman analysis.

## Methods

### Materials

Glyceryl trioctanoate (≥ 97%) was sourced from Sigma Aldrich, and octanoic acid (99%) was obtained from Thermo Fisher Scientific. Raman measurements of fatty acids were conducted using Corning™ 96-well half-area high-content imaging glass-bottom microplates. Lysogeny Broth (LB)-Miller medium, composed of 5 g/L yeast extract, 10 g/L sodium chloride, and 10 g/L tryptone, was purchased from Fisher BioReagents. M17 medium, composed of 2.5 g/L tryptone, 2.5 g/L peptone, 5 g/L soy peptone, 5 g/L meat extract, 5 g/L yeast extract, 19 g/L sodium glycerophosphate, 5 g/L lactose, 0.25 g/L magnesium sulfate, 0.5 g/L ascorbic acid was purchased from Fisher BioReagents. MRS medium, composed of 10 g/L proteose peptone, 10 g/L beef extract, 5 g/L yeast extract, 20 g/L dextrose, 2 g/L dipotassium phosphate, 5 g/L sodium acetate, 2 g/L ammonium citrate, 0.1 g/L magnesium sulfate, 0.05 g/L manganous sulfate, 1 g/L polysorbate 80 was purchased from Fisher BioReagents. YPD medium was composed of 20 g/L dextrose (Fisher BioReagents), 20 g/L peptone (BD) and 10 g/L yeast extract (MilliporeSigma).

### Sample preparation

The binary mixtures of glyceryl trioctanoate (GTO) and octanoic acid (OA) with different compositions were prepared by 1:1 serial dilution of OA using pure GTO, starting from 90% GTO and 10% OA (vol.%) in a 1 mL solution to an end composition of 99.98% GTO and 0.02% OA (vol.%) in a 1 mL solution. 150 µL of each sample was added to the microplate wells for Raman analysis. Samples were freshly prepared on three different days to evaluate inter-day variation.

### Cell culture

A single colony of *Escherichia coli* BL21(DE3) was picked from a Lysogeny Broth (LB)-Miller-agar plate and inoculated into a liquid LB-Miller medium. The cells were grown overnight at 37°C in a culture tube with constant shaking at 250 RPM. A single colony of *Saccharomyces cerevisiae* EBY100 was picked from a YPD-agar plate and inoculated into liquid YPD medium. Cells were cultured in a baffled flask at 30°C, 250 RPM for 24 h in YPD media. A single colony of *Lactococcus lactis subsp. cremoris (MG1363)* was picked from an M17-Agar plate and inoculated into a liquid M17 Medium with 0.5% Lactose. The cells were grown overnight at 30°C in a culture tube with constant shaking at 250 RPM. A single colony of *Lactobacillus reuteri (IDCC 3701)* was picked from a MRS-Agar plate and inoculated into a liquid MRS Medium. The cells were grown overnight at 37°C in a culture tube with constant shaking at 250 RPM.

The cell cultures were centrifuged to obtain a cell pellet, which was washed three times with 1 mL of phosphate-buffered saline (PBS). Subsequently, 10 μL of the washed cells were deposited onto a microscope slide, covered with aluminum foil, to form a thin film of intact cells. The film was allowed to dry completely before acquiring Raman spectra.

The Raman spectral data of engineered β-carotene producing *Saccharomyces cerevisiae* strain with reconstructed single, double, or triple mutations identified from adaptive laboratory evolution experiments analyzed in this work were acquired from prior publication ^21^. The spectra were collected using a Thermo Fisher Scientific DXR3 Raman microscope under identical measurement conditions, as described in this study. Information on the yeast strains is provided in Table S 14.

### Raman measurements

The majority of Raman spectra were collected with a Thermo Fisher Scientific DXR3 Raman microscope, designated as Instrument 1 (I1), using laser excitation at a wavelength of 780 nm and an output power of 20 mW. The instrument has an Olympus BX41 optical microscope and a thermoelectrically cooled charge-coupled detector (Thermo Fisher front-illuminated CCD system) with a 1024 × 256-pixel format, operating at −70 °C. The signal was calibrated using an internal polystyrene standard and a 10× objective (numerical aperture (NA) 0.25). The spot size was approximately 3.8 µm (Airy disk diameter, calculated as 1.22 λ/NA). For each sample, 50 spectra were collected with an exposure time of 15 seconds for two accumulations at different spots. 150 spectra were collected over three days (50 spectra for each sample per day). Similar settings were used to measure the Raman spectra of different cells, i.e., a laser excitation wavelength of 780 nm and an output power of 20 mW. A 50× (NA 0.5) objective was used. The spot size was approximately 1.6 µm(Airy disk diameter). Sixteen spectra were collected with an exposure time of 10 seconds for 12 accumulations at different spots for each sample. Forty-eight spectra were collected over three days (16 spectra for each sample per day).

The second Raman spectrometer (I2) was used to evaluate the influence of instrument variations. A custom-assembled Raman microscope system (Figure S 1), which integrates a commercial Raman spectrometer with a home-built microscope, instrument 2 (I2). The spectrometer is a portable iRaman Plus (BWTek, Plainsboro, NJ, USA), which features fiber delivery and utilizes 785-nm excitation light ^22^. The standard sampling probe at the output end of the fiber coupler was removed and replaced with a microscope adapter that connects to our modified microscope system. The 785-nm excitation light is focused to a flat-top spot with a diameter of 100 µm using a L Plan 20× (0.40 NA) long-working distance (12 mm) microscope objective. The light scattered from the sample travels back through the microscope objective to the fiber coupler, where it is separated from the laser light using a short-pass dichroic beam-splitter. This beam-splitter allows the 785 nm laser light to pass through while reflecting the Raman signal into the signal fiber of the Raman spectrometer. Data acquisition was performed using the BWSpec 4.0 software supplied with the iRaman Plus spectrometer. A schematic of the system is shown in the Figure S 1. Raman spectra were acquired at 20 different lateral positions within the liquid sample (all at a 2 mm depth below the liquid surface). The laser power was set to 10 percent, resulting in 40 mW measured after the 20× microscope objective. We set a laser exposure time of 15 s and two averaged spectra were taken at each measurement point. The same parameters were used for all samples measured with this system. Sample positioning was achieved using high-precision motorized sample translation stages (PLS-XY, Thorlabs Inc, Newton, NJ, USA), and focus adjustment was done with a high-precision vertical translation stage (ZFM2030, Thorlabs Inc). Both the sample and focus position can be adjusted with 1µm precision.

### Data processing of Raman spectra

We used the same methodology reported in the previous studies for the data analysis ^11, 23^. The spectra were first processed using asymmetric least squares (ALS) baseline correction for data processing. Then, the baselined spectra were vector normalized. The spectra were truncated in the range of 200-1800 cm^−1^. We truncated the spectrum to focus on the range with the highest signal quality and eliminate Raman silent region. Data processing was conducted with MATLAB 2023b.

### Instrument calibration

The instruments were first calibrated for wavenumber shift using a standard. This shift was adjusted to align the spectra from both instrument 1 (I1) and instrument 2 (I2). For intensity correction, we found the local maxima (peaks) where the intensity ratio between the spectra from I1 and I2 exceeded a threshold of 0.004.

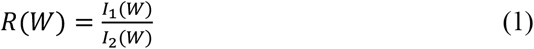

where R(W) is the ratio of the intensities of the two spectra I1 and I2 at wavenumber W.

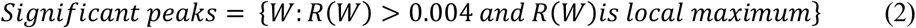

This threshold was selected to exclude noisy peaks, ensuring that only significant peaks were considered for correction and preventing overfitting. We used a third-degree polynomial to fit the identified peak locations and generated an intensity correction factor, achieving an R^2^ value of 0.8775 for the fit. This polynomial equation was then applied to correct the spectra collected from I2.

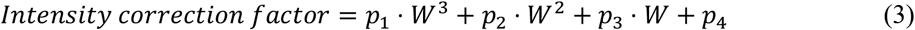

where *p*_1_, *p*_2_, *p*_3_ and *p*_4_ are coefficients and W is the wavenumber.

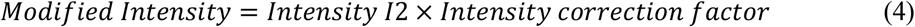

### Multivariate analysis and machine learning classification

We used a similar approach to multivariate analysis and machine learning classification, as discussed in previous studies ^10, 11, 23^. Before applying classifiers, the smoothed spectra were processed using multivariate statistical analysis to reduce complexity and extract the significant spectral features that explain the most variance. Discriminant analysis of principal components (DAPC) was used in this study. First, principal component analysis (PCA) was applied to reduce data complexity. Then, a supervised multivariate analysis, discriminant analysis, was used to further discriminate the dataset by correlating variation in the data with sample information. After feature extraction, standard machine learning classifiers were used to classify Raman spectra. In this study, we used Naïve Bayes (Gaussian), Support Vector Machine (SVM), K-Nearest Neighbors (KNN), and Neural Network (NN) to compare the performance of different algorithms.

We also used a Convolutional Neural Network (CNN) with PCA for feature extraction. The 1-D CNN model consists of five convolutional layers with 128, 64, 32, 16, and 8 filters, respectively, each followed by max-pooling to extract features from the input Raman spectroscopy data. The output is flattened and passed through two fully connected layers with 128 and 64 units, using ReLU activation and batch normalization. The final SoftMax layer predicts class probabilities. The model was compiled with the Adam optimizer and sparse categorical cross-entropy loss, optimizing for accuracy.

To assess the suitability of the classification algorithm, a 5-fold cross-validation was performed. In brief, 5 different training and validation sets were established by randomly selecting from the Raman spectra data.

### Establishing spectral library with artificial noises

To account for the impact of noise on classification accuracy, we generated the simulated spectral data of the mixture of GTO and OA. The spectra of lipid mixtures were created using weighted blending, which calculates the weighted average of the spectra of pure GTO and pure OA ^24, 25^ The mathematical equation is stated below:

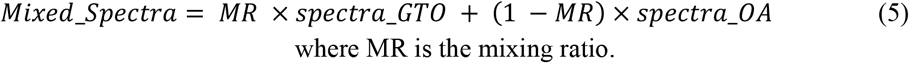

We used a Gaussian noise model to introduce noise to the simulated spectral data ^26–30^. The mathematical expression for the model used is stated below:

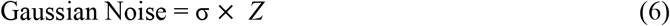

where *Z* is a random variable that follows a standard normal distribution, *Z*~𝒩(0,1), i.e., mean 0 and variance 1, and σ is the standard deviation (noise level).

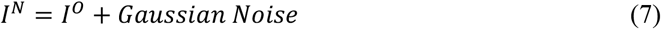

where *I*^*O*^ is the original intensity, and *I*^*N*^ is the new intensity with noise.

Machine learning and deep learning algorithms depend heavily on large datasets. To address this, we generated 1,000 spectra with random noise. In this study, we varied σ across a broad range (0.5-60) to evaluate how increasing noise levels affect classification accuracy.

### Spectral similarity calculations

To evaluate the similarity of Raman spectra, we used established metrics from the literature, such as the Pearson correlation coefficient and the Pair (Chebyshev distance) ^31, 32^.

The mathematical expression for each metric is stated below:

Let *A* and *B* be two spectra, where each spectrum is represented as a vector;

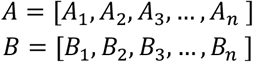

For these spectra:

- ***Pearson correlation coefficient (PC)***

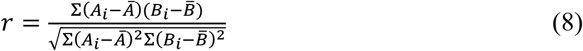

Here, *A*_*i*_ and *B*_*i*_ are intensity values at the *i*-th wavelength, and 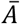 and 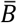 are the mean intensities in the spectrum. Like cosine and first difference cosine similarity, the Pearson correlation coefficient is 1 for a perfect positive correlation between the spectra, 0 for no correlation, and −1 for a perfect negative correlation between the spectra.
- ***Pair distance (PD)***

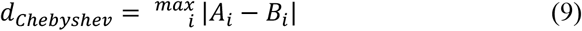

Here, *A*_*i*_ and *B*_*i*_ are intensity values at *i*-th wavelength For identical spectra, the pair distance is 0, and it goes to ∞ for dissimilar spectra, depending on how dissimilar the spectra are.

## Results and Discussion

### Evaluating impact of spectral noise and similarity using simulated spectra

To evaluate the impacts of spectral similarity on classification accuracy, a serial dilution of octanoic acid (OA) mixed with glyceryl trioctanoate (GTO) were prepared. GTO is a triglyceride of OA containing medium-chain (C8) saturated fatty acid chains. These lipids serve various functions, including acting as an antibacterial agent, a human metabolite, and a metabolite in *Escherichia coli* ^33, 34^. Due to their similar chemical structures, GTO and OA exhibit comparable Raman spectra (Figure 1). The major spectral differences are at 1662 cm^−1^ and 1746 cm^−1^, corresponding to the carboxylic group in OA and the aliphatic ester in GTO, respectively. Other peaks in the region 800-1470 cm^−1^ are contributed by the fatty acid chain expansion of n-alkanes ^10, 35^.

**Figure 1.**
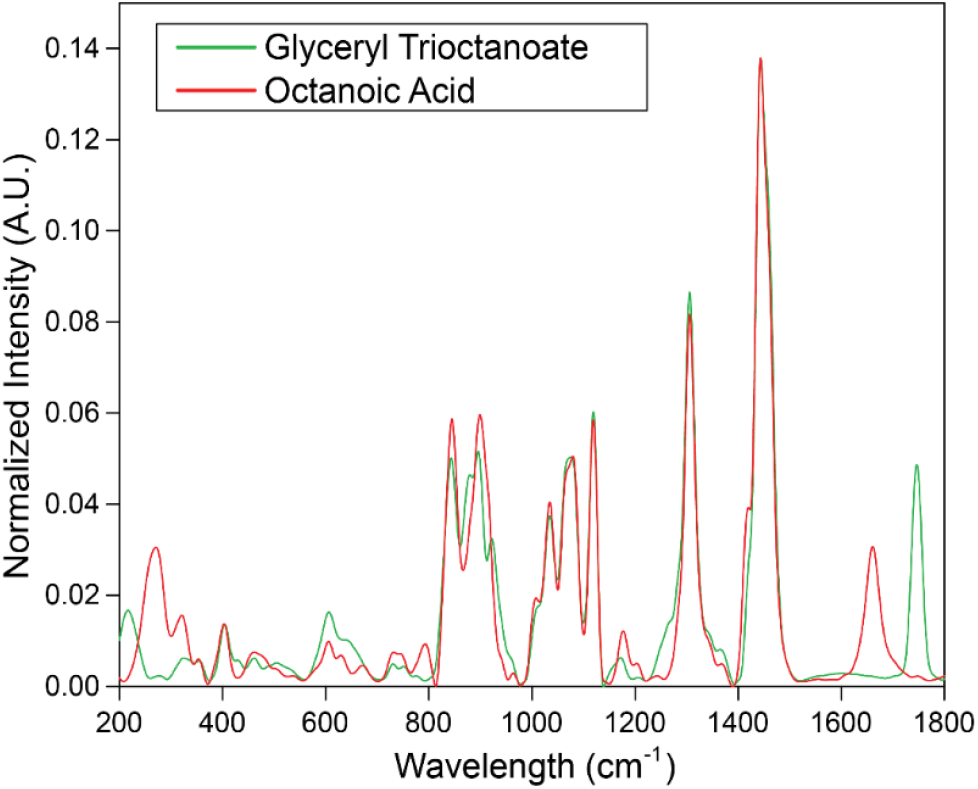
Average normalized Raman spectra (total number of spectra for each component (n)= 50) of pure Glyceryl Trioctanoate (GTO) and pure Octanoic Acid (OA).

Prior studies have shown that the spectrum of a mixed sample should closely resemble a linear combination of the spectra of its individual components ^4, 36^. The spectra of GTO and OA mixture can be simulated by equation 5 ^24, 25^. We began with a mixture composition of 25 vol.% GTO and 75 vol.% OA, and progressively reduced the OA concentration, reaching as low as 99.98 vol.% GTO and 0.02 vol.% OA in the mixture. This allowed us to create a dataset with varying degrees of similarity between the tested samples. Two different similarity metrics that were widely used in the literature are used to determine the similarity between the spectra, including Pearson correlation coefficient (PC) and Pair (Chebyshev) distance (PD) ^31, 32^. The spectral similarity scores between pure GTO and OA: PC=0.9151 and PD=0.0457. The compositions and the simulated Raman spectra of the mixture samples are shown in Table S1 and Figure S2. We used the 100% GTO as the reference, and we calculated the similarity scores between the spectra of the reference and other compositions (Table S2). As the composition difference decreases, i.e., as the OA content in the mixture is reduced, the spectral similarity increases.

In addition to spectral similarity, spectral variations could influence the classification accuracy. The variations could be contributed by various factors, such as sample preparation, natural variations in biological samples, and inconsistencies among different instruments, laboratories, or technicians. To introduce random noise into our simulated spectra, a Gaussian noise model was employed, allowing us to account for the impact of spectral noise on classification accuracy ^26–29^. Table S3 presents the average standard deviation of the spectra at each noise level.

To evaluate how spectral noise influences spectral similarity, we compared similarity scores between individual spectra within the same group and between two compositions. Figure 2 presents a scatter plot of the similarity scores, using PC, at noise levels (σ) of 0.5 and 15. At low noise (σ = 0.5), the individual spectra within the same composition group (intra-group) have a consistent spectral pattern, showing a high similarity. The similarity score of inter-group (between two compositions) is significantly lower than intra-group similarity. However, as noise increases (σ = 15), the values of intra-group similarity significantly spread out, and the score is at the same level of the inter-group comparison, which will lead to the reduction of classification accuracy. The results indicate that the similarity score depends on the likeness of spectral profiles and is heavily influenced by the noise levels in the spectra ^31^. PD yielded comparable results (Figure S 3).

**Figure 2.**
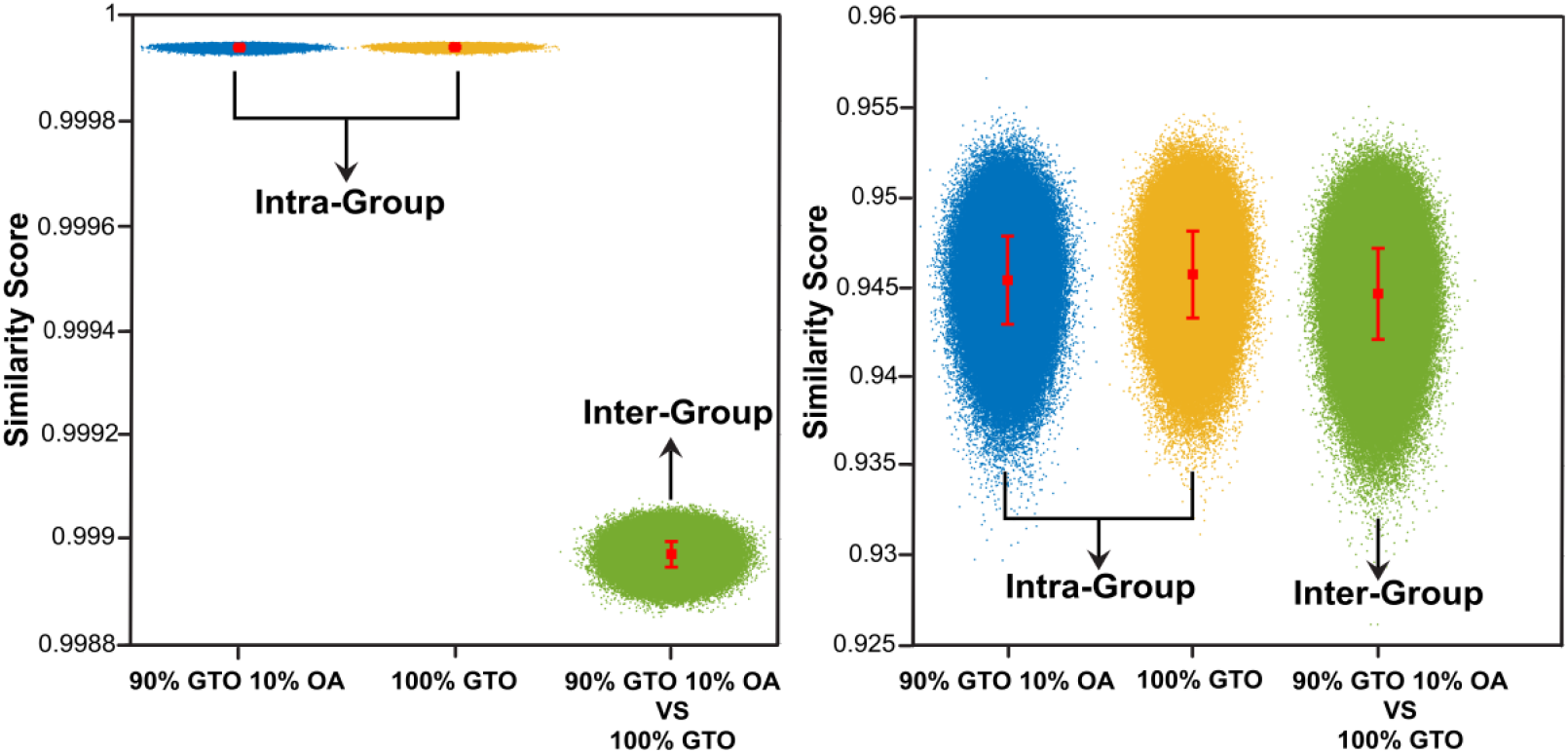
Scatter plot for PC similarity score for 90% GTO 10% OA compared to 100% GTO for noise levels (σ) (a)0.5 and (b) 15. As noise increases (σ = 15), intra-group similarity scores become substantially more spread out and overlap with inter-group scores, resulting in reduced classification accuracy. Red dots and error bars represent the mean and standard deviation of similarity scores for each group.

### Impact on classification accuracy

To evaluate the performance of different ML algorithms, we employed several widely used machine learning algorithms: Naïve Bayes (Gaussian), Support Vector Machine with a linear kernel (SVM-Linear), K-Nearest Neighbors with Euclidean distance (KNN-Euclidean), Neural Network (NN), and Convolutional Neural Network (CNN) ^10, 11, 23^. We applied DAPC (Discriminant Analysis of Principal Components) for dimensionality reduction for the models Naïve Bayes (Gaussian), SVM-Linear, KNN-Euclidean, and NN. For CNN, the spectral data was imported directly without dimensionality reduction.

To evaluate the effect of noise on classification accuracy, we applied all six machine learning models for binary classification, distinguishing the reference mixture (99.9805% GTO, 0.0195% OA) spectra from those of other mixtures. Figure 3 and Table S6 report the accuracy scores for binary classification of various mixtures using the SVM-Linear model. As expected, classification accuracy decreases with increasing noise levels in the spectra. Binary classification results for all models are detailed in Tables S4–S9. All models exhibit comparable performance, with misclassifications increasing at higher noise levels, resulting in a greater standard deviation in the spectra, even when compositional differences are substantial.

**Figure 3.**
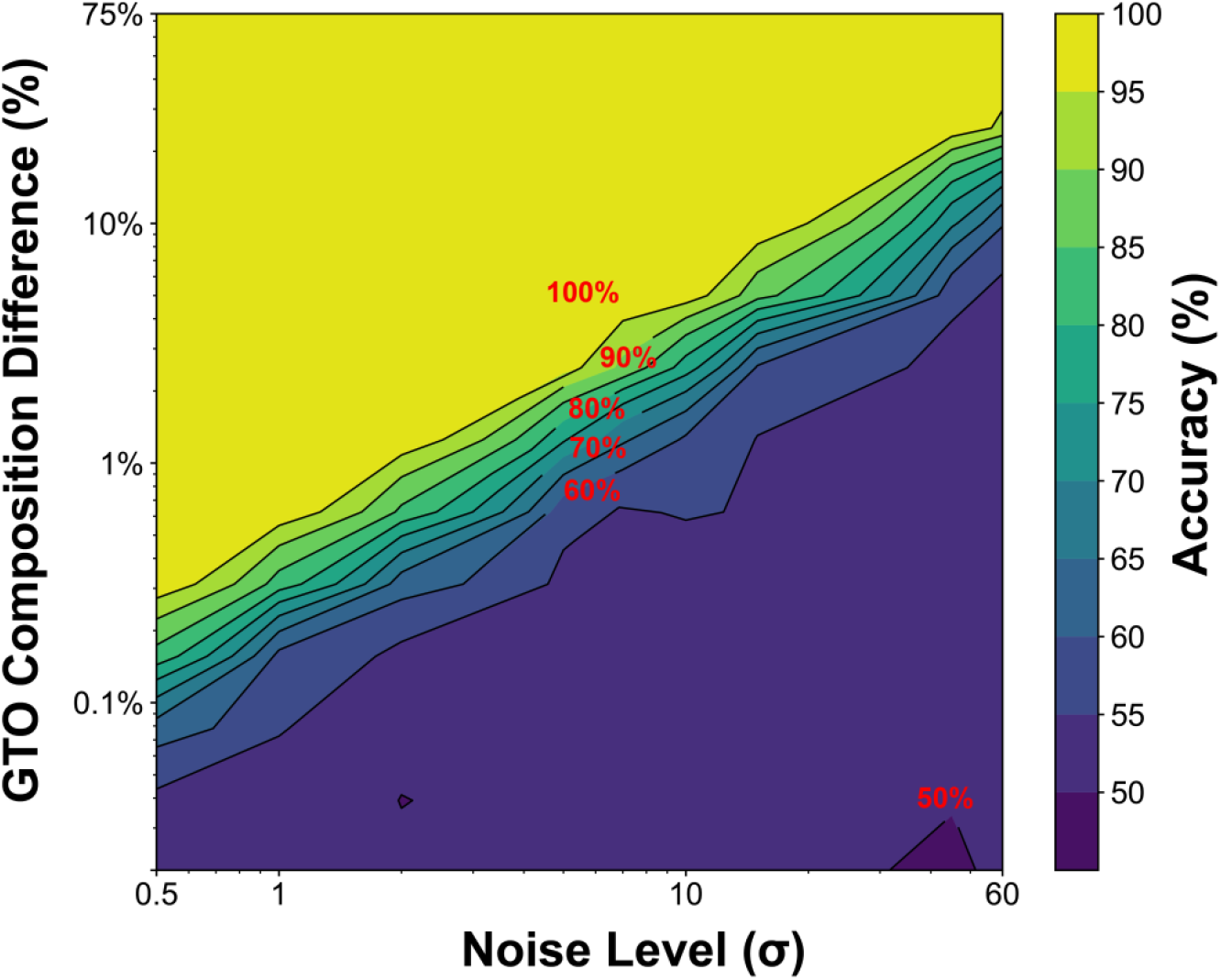
Contour Plot for Binary Classification Accuracy: Illustrating the relationship between GTO Composition Difference (%) and Noise Level (σ), with color gradients representing accuracy percentages using SVM-Linear.

Increasing the number of classes increases the complexity of machine learning classifiers, thereby reducing classification accuracy. ^37^. In addition to binary classification, multiclass classification was performed on the simulated data. Figure 4 shows the row-wise normalized confusion matrices. When the noise level is σ = 0.5, the SVM model is capable of distinguishing samples differing by more than 0.605 vol% in composition with accuracy exceeding 99% (Figure 4a). At σ = 5, however, multiclass classification requires composition differences larger than 5 vol% (Figure 4b). Figures S4-S12 show confusion matrices for multiclass classification of simulated data with different levels of noise using SVM (Table S10 includes confusion matrix label information). The results indicate that the noise levels in the spectra heavily influence classification accuracy.

**Figure 4.**
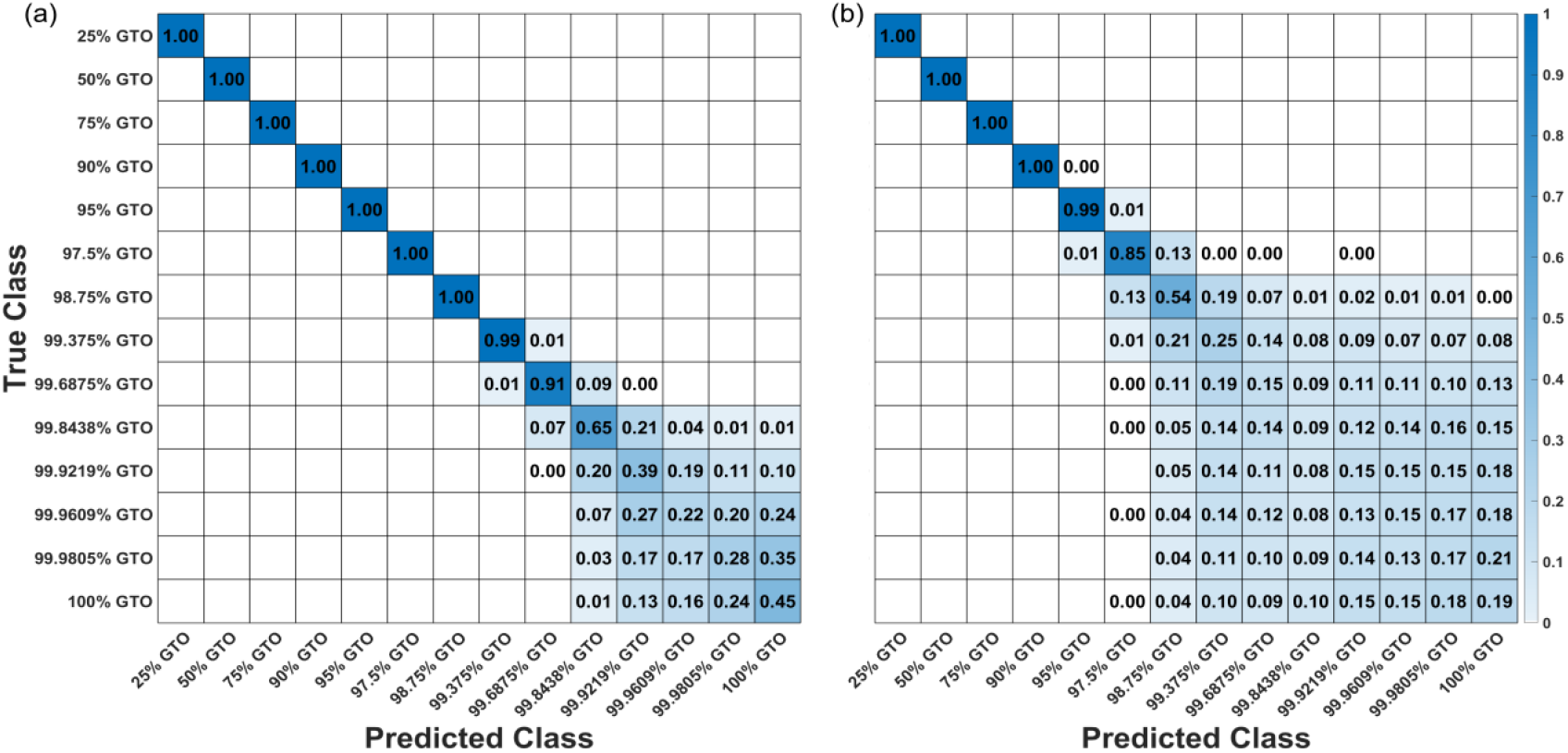
Row-wise normalized confusion matrix for multiclass classification of simulated mixtures for different noise levels, (a) σ=0.5 (average standard deviation 1.67×10-4) and (b) σ=5 (average standard deviation 1.62×10-3), using SVM-Linear.

### The influence of experimental factors on classification accuracy

Experimental factors such as sample preparation and various other sources contribute to the noise in the spectra, including environmental factors such as room lighting, background signals, and fluorescence. Instrumental noise includes dark current, photon shot, and read noise, impacting the overall spectral data quality. We prepared mixtures with the same compositions of GTO and OA to experimentally validate the results obtained from simulated data.

To assess the impact of inter-day sample preparation on the standard deviation between spectra, samples were freshly prepared on three different days to evaluate inter-day variation. For each sample, fifty spectra were collected on each day. Table S11 shows the average standard deviation values for inter-day and intra-day sample preparations. The mean standard deviation for inter-day variation is 4.69×10^−4^ (corresponds to noise level (σ) ~1.45), and for intra-day variation, it is 4.10×10^−4^ (corresponds to noise level (σ) ~1.25). The lower average standard deviation in intra-day measurements compared to inter-day sample preparation suggests inconsistencies in the sample preparation process across different days.

A similar binary classification was performed first, as discussed in the above section. The classification results for different composition differences are presented in Figure S13. We observe a similar trend in accuracy scores for both intra-day and inter-day sample preparation. As the composition difference narrows, the accuracy score tends to decrease. The classification accuracy for inter-day and intra-day data drops below 95% as the composition difference reaches 0.605 vol. % (corresponds to 1.85 mol%). We observed similar results for all the different machine-learning models. While the composition difference becomes smaller, the similarity between the spectra increases, resulting in more misclassifications.

When we compare the results of simulated and experimental data with similar standard deviation values, we observe that the classification accuracy is comparable for both data sets at similar noise levels in the spectra (Figure 5 and Tables S12 and S13).

**Figure 5.**
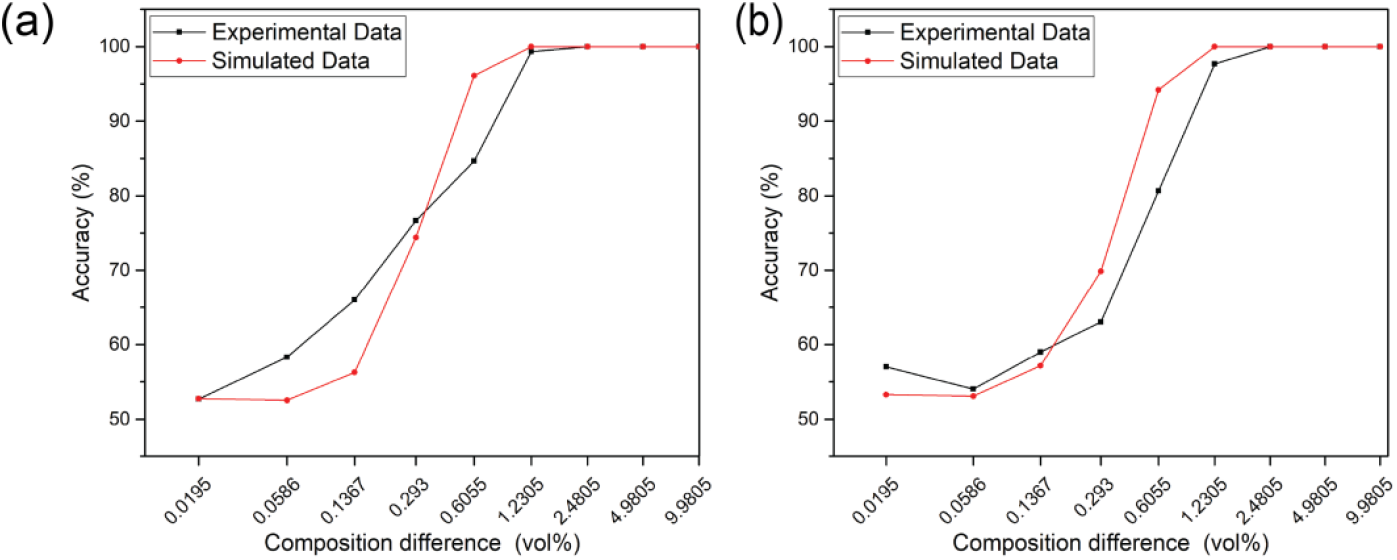
Comparison between experimental and simulated results with similar standard deviation values,(a) Intra-day and (b) Inter-day, using SVM-linear.

We conducted a similarity comparison and multiclass classification on the experimental data using the same six models, employing an approach similar to the one described previously. The sensitivity reflects an ML model’s ability to accurately identify an actual composition. For both intra-day and inter-day classification, the SVM model is capable of distinguishing samples differing by more than 1.25 vol.% (approximately 3.78 mol%) in composition with accuracy exceeding 95% (Figure 6a and 6b, respectively). Figures S14-S23 show confusion matrices using the other models. Similar to simulated data, all the models have comparable classification performance.

**Figure 6.**
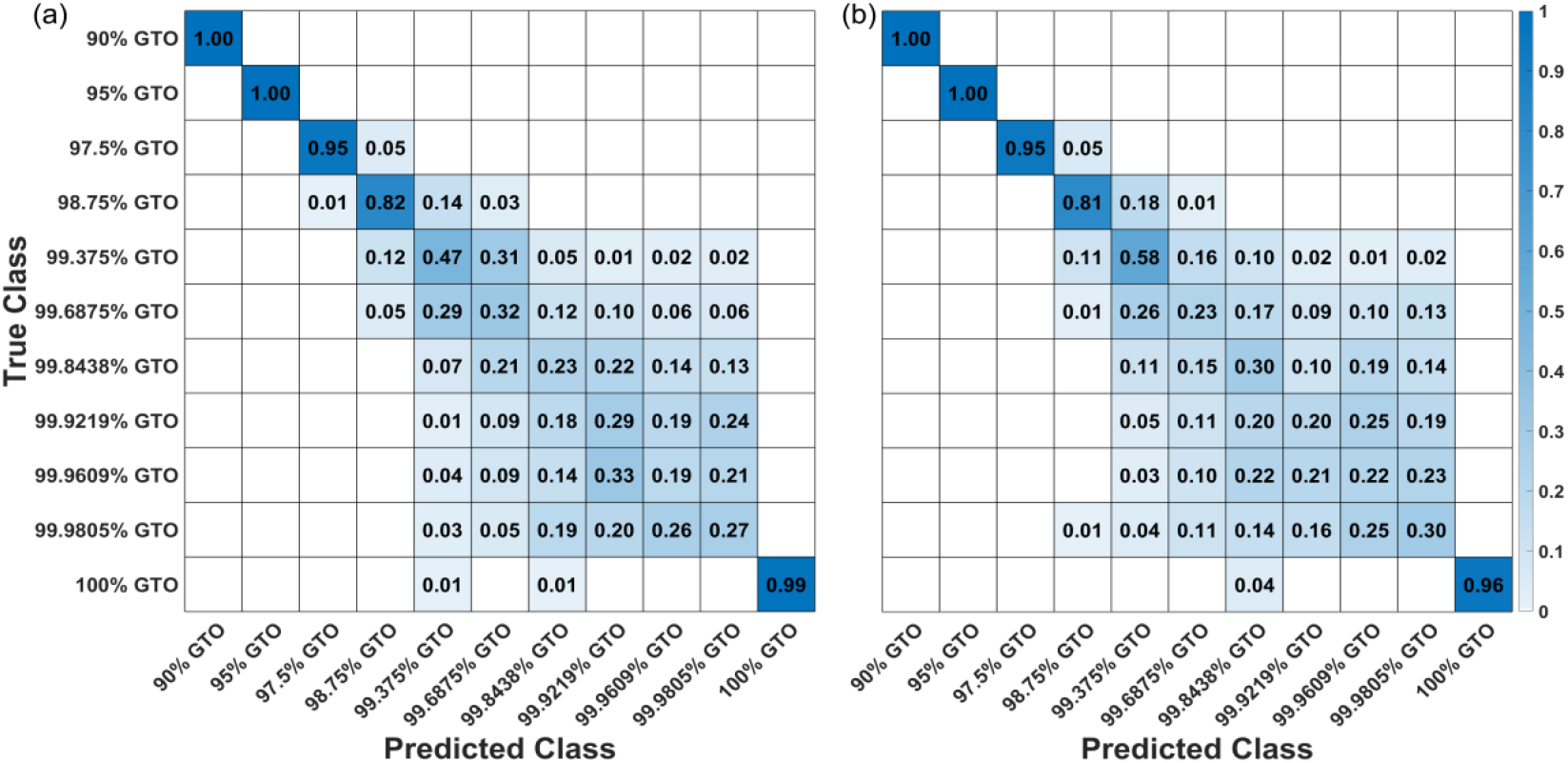
Row-wise normalized confusion matrices for multiclass classification of (a) intra-day data and (b) inter-day data using SVM-Linear.

As previously discussed, spectral noise significantly affects classification performance. One effective way to improve detection accuracy is to increase the SNR. Averaging multiple Raman spectra progressively enhances the SNR, resulting in more precise and reliable spectral features ^38^. In this study, every five consecutive spectra were averaged (n = 5). This averaging reduced the mean standard deviation to 2.75 × 10^−4^ for inter-day measurements and 1.91 × 10^−4^ for intra-day measurements. With averaged spectra, the SVM model achieved >90% classification accuracy for both intra-day and inter-day datasets when distinguishing samples with composition differences greater than 0.625 vol.% (Figure 7). Overall, spectral averaging substantially reduced variability and significantly improved classification performance. Therefore, averaging multiple spectra is a simple yet effective preprocessing technique for minimizing noise-induced variability and enhancing classification accuracy in Raman spectral analysis.

**Figure 7.**
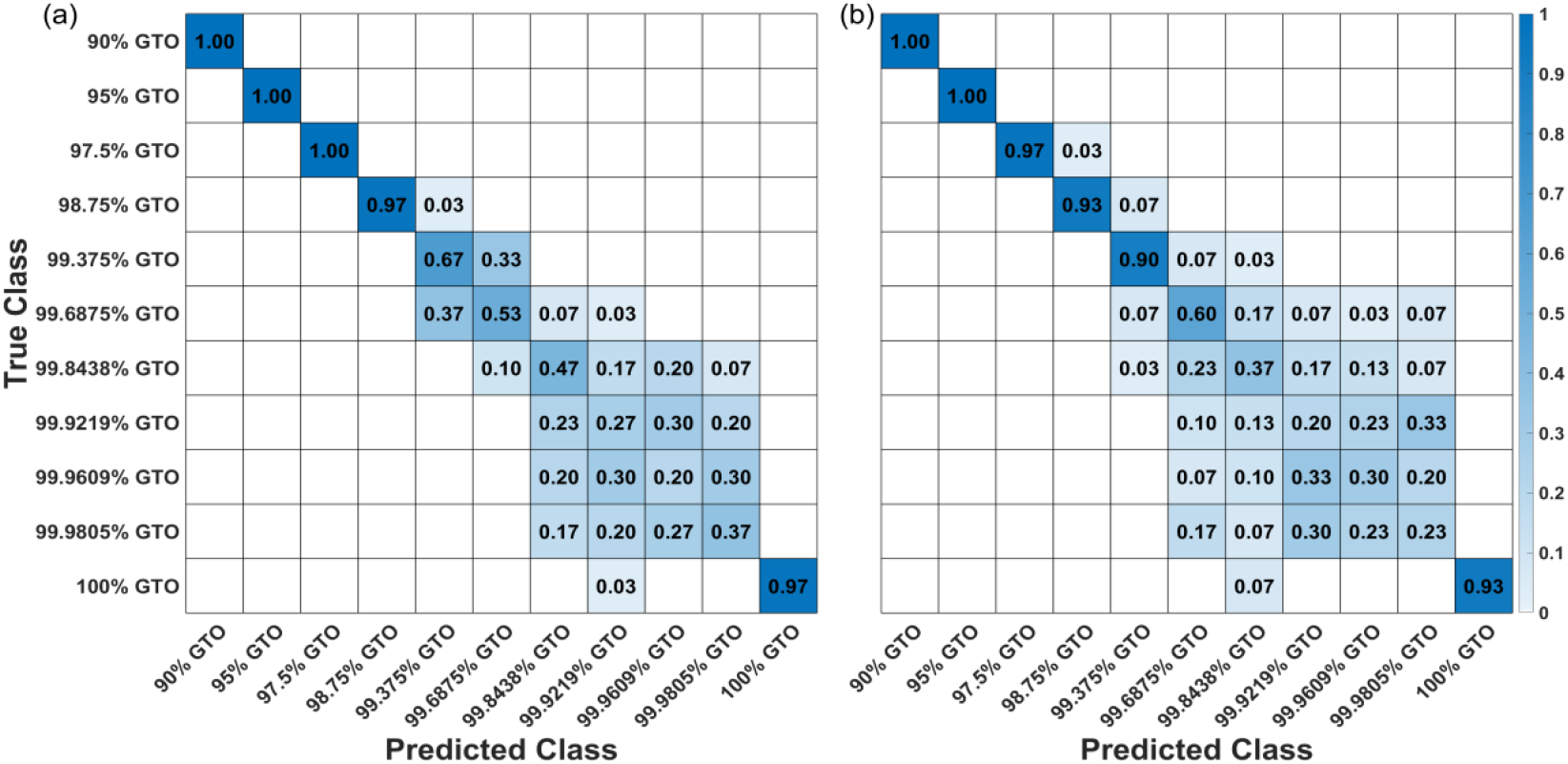
Row-wise normalized confusion matrices for multiclass classification of averaged spectra (every five consecutive spectra were averaged) using SVM-Linear (a) intra-day data and (b) inter-day data. Spectral averaging substantially reduced variability and significantly improved classification performance.

### Transfer learning across different spectrometers

When spectra are collected from different Raman spectroscopes in varying environmental conditions (such as different labs), inconsistencies can occur in the spectra, even for identical samples. Variations in the quantum efficiency of the CCD, laser wavelength, grating efficiency, reflection or transmission efficiencies of optical components, sampling geometry, spectrometer resolution, power applied to the sample, and spectral response can significantly impact the Raman spectra obtained from each instrument ^39, 40^. These variations introduce an additional layer of complexity to the data, impacting ML performance. Figure7a shows spectra for samples collected from two Raman spectrometers, a Thermo Fisher Scientific DXR3 Raman microscope (I1) and a custom-assembled Raman microscope system (I2), which uses a portable iRaman Plus (BWTek, Plainsboro, NJ, USA).

By comparing the intensities of multiple Raman bands at 606, 896, 1306, and 1444 cm^−1^ (Figure 8a), we found that spectra acquired on instrument I2 require intensity correction. Several established calibration methods have previously been employed using NIST standards and have proven effective ^41^. For example, Jason et al. used NIST SRM-2241 to derive an instrument-specific correction function by dividing the certified reference spectrum by the measured SRM-2241 spectrum; the corrected sample spectrum was then obtained by multiplying the raw Raman spectrum by this correction function ^39^. Similarly, Mark et al. employed a standard white-light source to characterize spectrometer throughput and collection efficiency, generating an instrument response function that was subsequently applied to correct recorded spectra ^42^.

**Figure 8.**
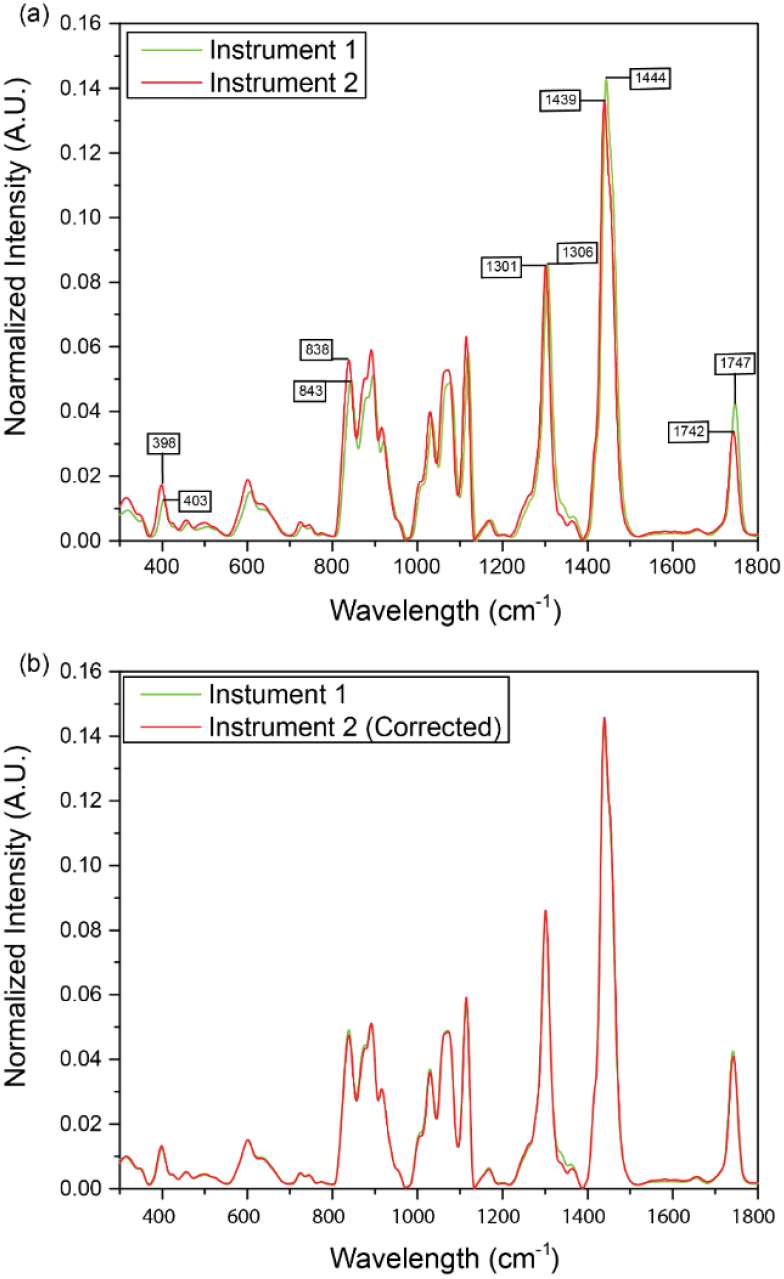
Raman Spectra for 90% GTO 10% OA collected from two different instruments. (a) Raman spectra before correction and (b) Raman spectra after correction.

We used a similar approach discussed in the published work ^39^. First, we calculated the intensity ratio between the spectra collected from the two instruments. Then, we applied a threshold, considering only peaks where the ratio exceeded a specific value for further correction. This approach helped us to isolate and highlight the most relevant spectral peaks. Based on these criteria, we filtered the significant peaks for intensity correction. We then used a 3rd-order polynomial to fit the selected peaks (to avoid overfitting), generating a polynomial function for intensity correction. Figure 8b shows spectra after shift and intensity correction.

We employed transfer learning by training the machine learning model on data acquired from instrument I1 and evaluating its performance on data from instrument I2. The classification results (Figure 9) show that transfer learning can successfully classify spectra across instruments.

**Figure 9.**
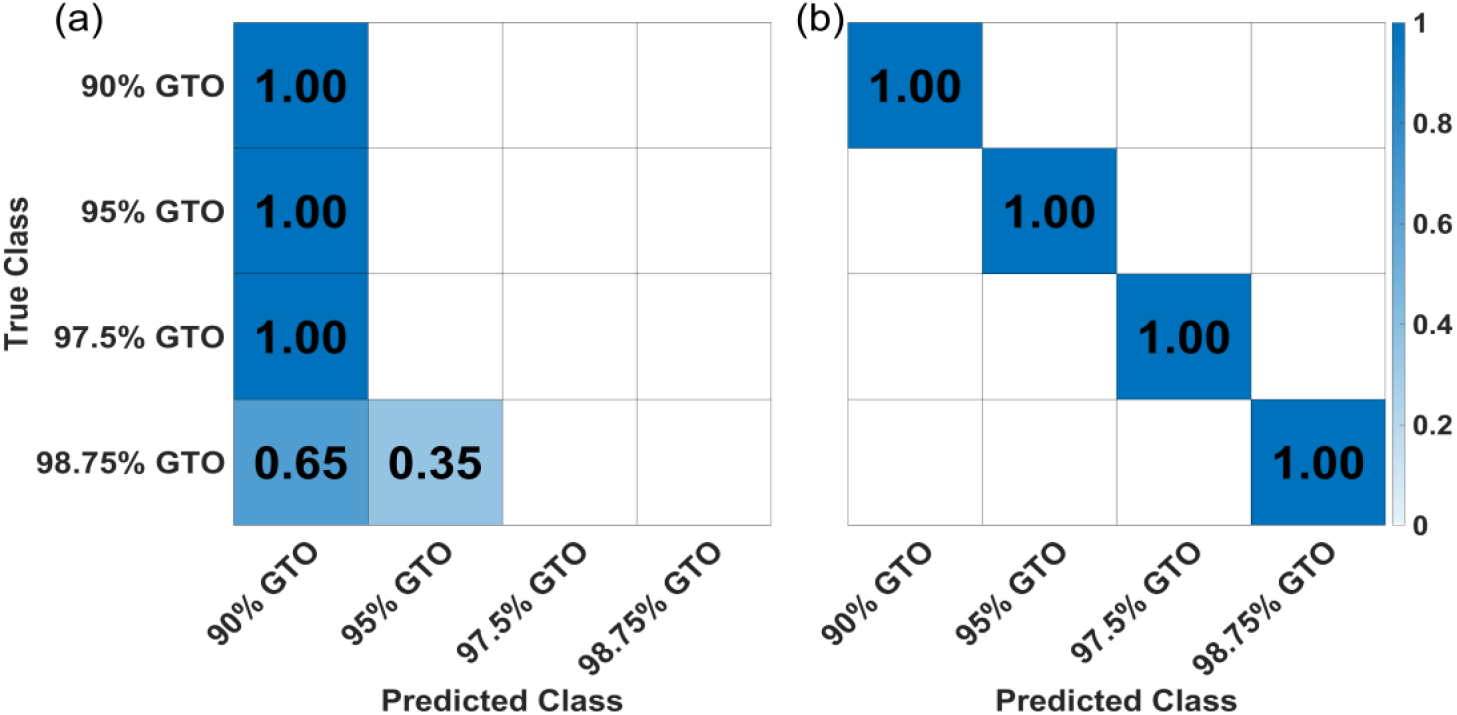
Row-wise normalized confusion matrix for inter-instrument classification using SVM (a) without spectral correction and (b) with spectral correction.

### Limiting factors in single-cell analysis

Machine learning-assisted Raman spectroscopy has been widely used to classify different types of cells at the single-cell level, such as bacteria, and their metabolic activities ^43–45^. It has also been used to classify various cancerous and non-cancerous cells, including breast cancer diagnosis, lung cancer, and esophageal squamous cell carcinoma ^46, 47^. However, cell-to-cell variations are observed among individual cells in a population, even when they are genetically identical and in the same environment. As a result, the intrinsic cell-to-cell variation can be challenging to extract meaningful information from the complex Raman spectra ^48^.

Bioengineering and metabolic engineering have made it possible to engineer cells to manipulate cellular processes, with the possibility of single-gene editing ^21, 49, 50^. We tested the ability of ML-assisted Raman spectroscopy to classify these mutations. Here, we re-analyzed the Raman spectral data from previously published work ^21^. Potential beneficial mutations for β-carotene production were identified from hyperproducers isolated from adaptive laboratory evolution experiments of an engineered *Saccharomyces cerevisiae* strain YLH2; Godara et al. studied the impact of these mutations on β-carotene yield via site-directed mutagenesis of the β-carotenogenic strain YLH2. In this study, Raman spectra from yeast strains containing single, double, and triple mutations were collected, and machine learning was employed to classify the cells. The labels for single mutations are YAG01, YAG02, YAG03, YAG04, YAG05, YAG06, YAG07, YAG08, YAG09, and YAG10; for double mutations, they are YAG17, YAG20, and YAG22; and for triple mutations, YAG23 and YAG28. We also collected the Raman spectral data of different bacteria, including *Escherichia coli (E. coli), Lactococcus lactis (L. lactis), Lactobacillus reuteri (L. reuteri)*, and *S. cerevisiae (EBY100)*. The information on the selected microorganisms is shown in Table S14. Figure S24 includes the averaged Raman spectrum for each class of microorganisms.

Due to inherent cell-to-cell heterogeneity, the spectral variations in yeast and bacterial cells are substantially larger than those observed in the well-controlled GTO-OA mixtures. The average standard deviations range from 3.46×10^−3^ to 1.01×10^−2^, which are approximately two orders of magnitude higher than those of the GTO-OA system (Table S15). Such large spectral variations are expected to adversely affect classification accuracy. We first evaluated binary classification performance (Figure 10 and Figures S25-S29). Despite the high spectral variability, the model achieved 100% accuracy in distinguishing the YLH2 strain from the bacteria (*E. coli, L. lactis*, and *L. reuteri*) and the wild-type *S. cerevisiae* strain (EBY100), owing to their relatively low spectral similarity. For the genetically engineered strains, although spectral similarity was generally higher, many were still classified with high accuracy. Specifically, the following strains achieved near-perfect binary classification: single mutants YAG01, YAG02, YAG03, YAG04, YAG06, YAG09, and YAG10; double mutants YAG17 and YAG20; and the triple mutant YAG28. However, classification accuracy dropped below 95% for YAG05, YAG07, YAG08, and YAG23.

**Figure 10.**
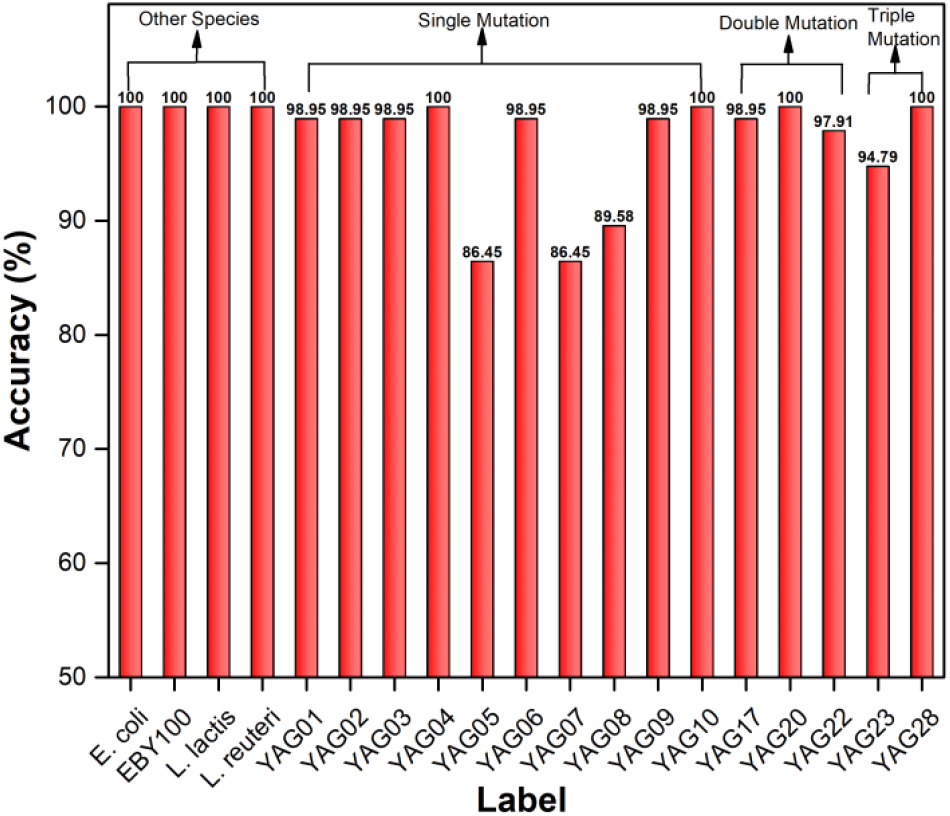
Binary classification results for all the cells compared to YLH2 using SVM-Linear.

We also performed multiclass classification (Figure 11 and Figures S30-S34). *E. coli*, EBY100, *L. lactis*, and *L. reuteri* were identified with high accuracy. In contrast, misclassification rates increased substantially for the mutated *S. cerevisiae* strains. Owing to extensive cell-to-cell heterogeneity and high spectral similarity among closely related genotypes, ML-assisted Raman spectroscopy was unable to reliably distinguish genetically similar *S. cerevisiae* strains at the single-cell level.

**Figure 11.**
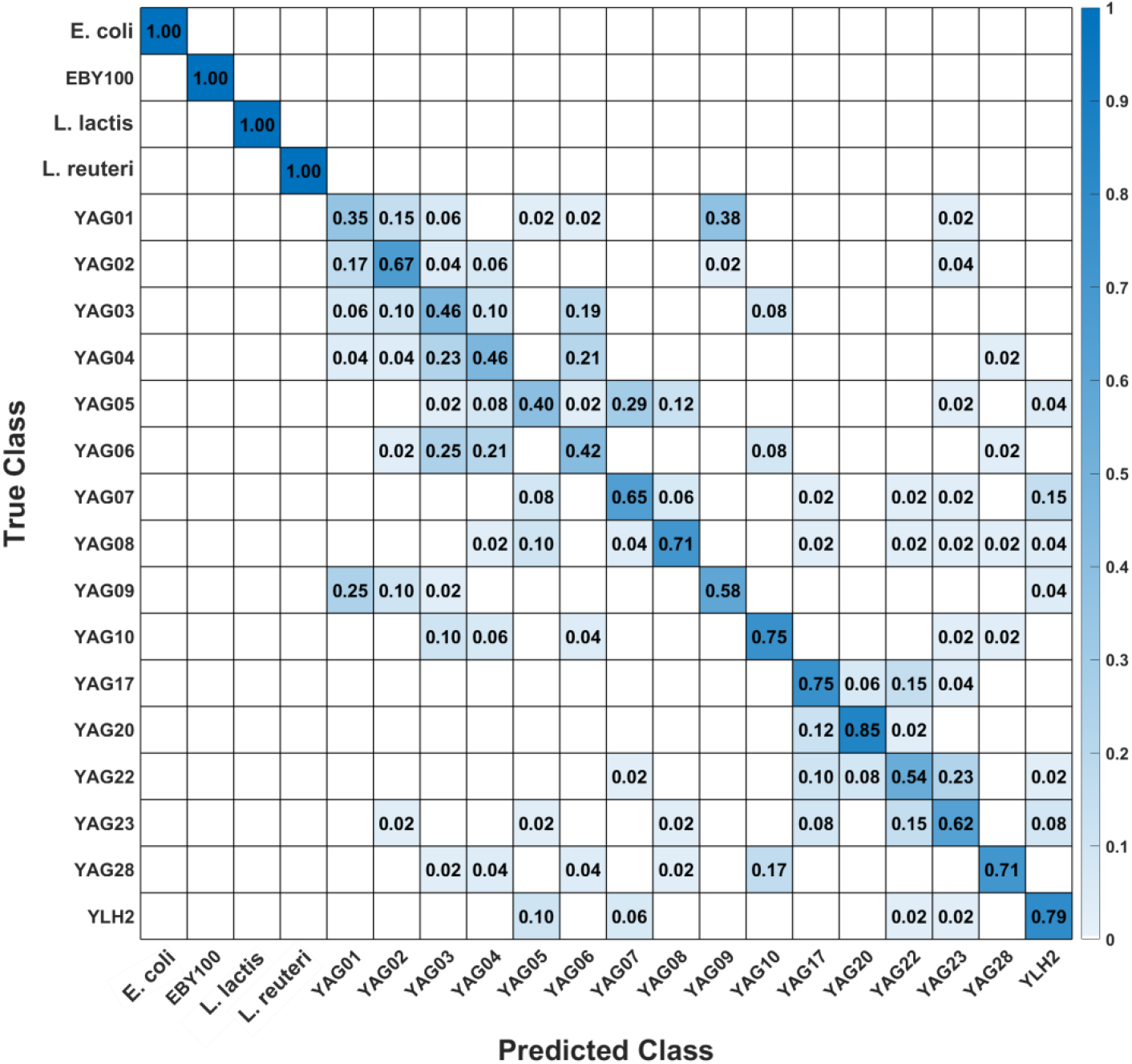
Row-wise normalized confusion matrix for multiclass classification of cells using SVM.

As previously described, averaging multiple spectra reduces random noise and variability, thereby decreasing the standard deviation across measurements and potentially improving classification accuracy. To evaluate this effect, we averaged the spectra acquired from individual microorganisms (n = 8 per class). The resulting confusion matrix for cell classification is shown in Figure 12. Overall, spectral averaging led to markedly improved classification performance compared to the use of single-cell spectra. In particular, classes that were frequently misclassified without averaging, such as YAG20 and YAG23, achieved 100% classification accuracy when spectra were averaged. Similarly, substantial gains in accuracy were observed for classes YAG02, YAG03, and YAG17 following spectral averaging.

**Figure 12.**
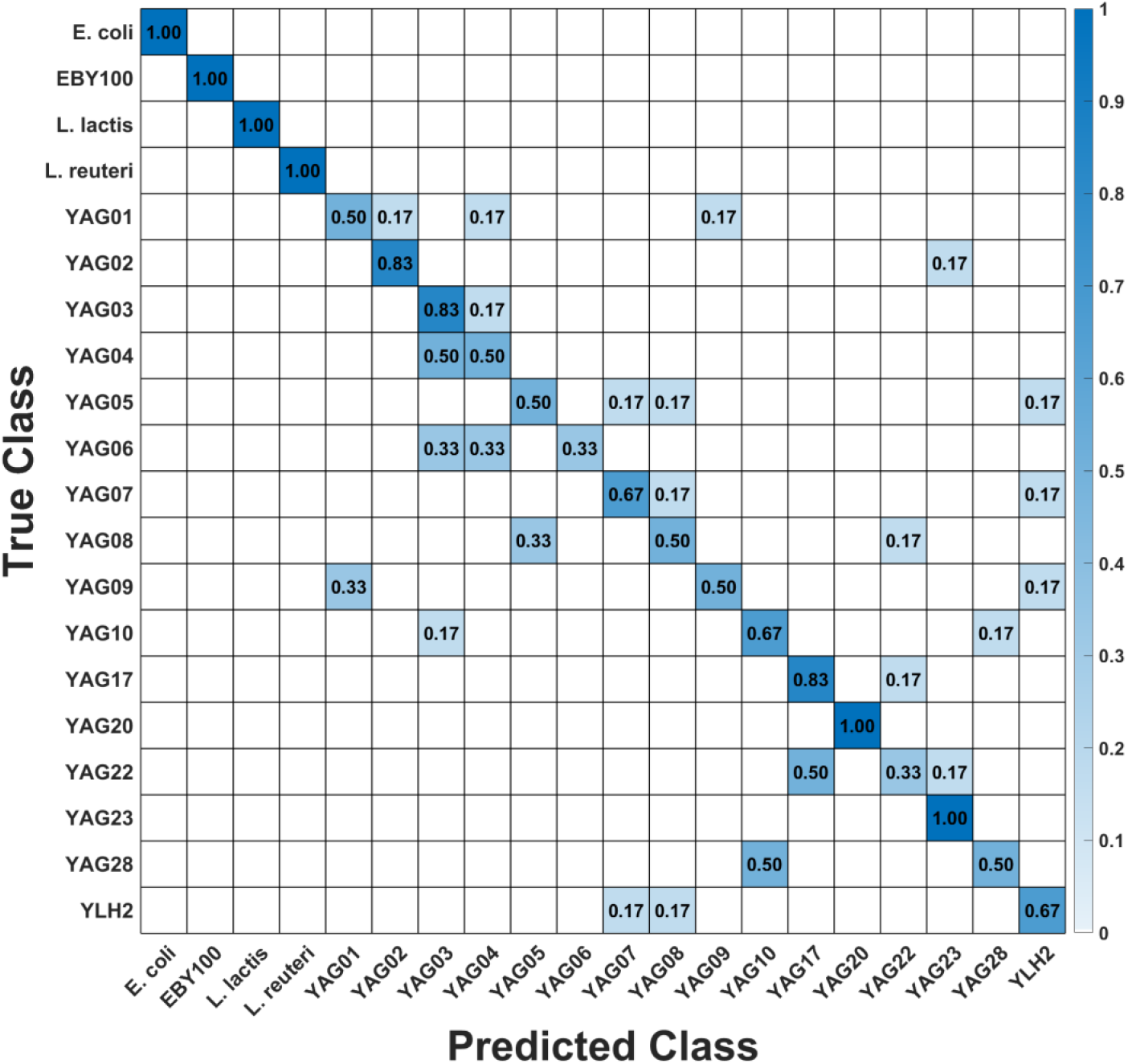
Row-wise normalized confusion matrix for multiclass classification of cells using SVM-Linear.

## Conclusions

Machine learning (ML)-assisted Raman spectroscopy is a powerful tool in analytical science. However, its performance is strongly influenced by spectral variations and the high spectral similarity between samples. In well-controlled binary mixtures of glyceryl trioleate and oleic acid (GTO-OA) with low noise levels, we successfully classified samples differing by as little as 1.85 mol% in composition. Further improvement in classifying highly similar samples can be achieved by reducing spectral variation through averaging multiple measurements. Careful experimental design can minimize both intrinsic noise sources (e.g., dark current, photon shot noise, and readout noise) and extrinsic variations (e.g., inconsistencies in sample preparation, ambient lighting, background signals, and fluorescence). Moreover, proper instrument calibration enables successful transfer learning, allowing a universal ML model trained on one Raman spectrometer to be effectively applied to others.

ML-assisted Raman spectroscopy has also been employed for single-cell classification of various cell types, including microorganisms and cancer cells ^43–47^. However, biological samples are inherently complex and diverse, exhibiting a wide range of Raman-active vibrational modes arising from biochemical components such as lipids, proteins, nucleic acids, and carbohydrates^1, 51^. Significant cell-to-cell spectral variation exists even among genetically identical cells grown under identical conditions. This heterogeneity arises from both intracellular randomness and external factors related to population dynamics ^52–54^, posing substantial challenges for single-cell analysis using Raman spectroscopy.

Our results demonstrate that ML-assisted Raman spectroscopy can successfully distinguish different microorganisms at the single-cell level despite high spectral variability. However, it failed to reliably differentiate *S. cerevisiae* strains carrying single, double, or triple mutations. Since Raman spectroscopy primarily reflects phenotypic differences—particularly in Raman-active molecules such as unsaturated lipids—greater genetic modifications that induce more pronounced biochemical alterations would increase spectral differences and improve classification accuracy. For highly similar engineered *S. cerevisiae* strains, reducing spectral variation is critical to achieve successful classification. Reduction of spectral variations may be effectively achieved by averaging Raman spectra acquired from a population of cells rather than relying on individual single-cell measurements.

In summary, the main limitations in ML-assisted Raman spectroscopy are data quality and spectral similarity. To achieve robust and reliable classification results, it is essential to pay close attention to sample preparation, data acquisition protocols, measurement conditions, and instrument calibration.

## Author Information

### Author Contributions

A.Y and H.W. contributed to the study design. A.Y. performed the experiments. A.F., A.J.V., and M.C. helped to perform experiments on the home-built Raman microscope. S.G. and S.K. provided cells. A.Y., A.B., N.A., and A.A. conducted data analysis. A.Y. and H.W. contributed to data interpretation and drafted the manuscript. Q.S., K.K., and X.Y. reviewed and edited the manuscript. All authors reviewed the manuscript.

### Notes

The authors declare no competing financial interest.

## Acknowledgements

The authors gratefully acknowledge the invaluable support from multiple organizations, including National Science Foundation (2114203 to H.W., 2114188 to K.K. & 2514387 to Q.S.), National Institute of General Medical Sciences (R35GM156609 to H.W.), Air Force Office of Scientific Research (FA9550-20-1-0366 to Z.Y.), and Robert A. Welch Foundation (A-1261 to Z.Y.). The home-built Raman microscope system is based upon work supported by the U.S. Department of Energy, Office of Science, Office of Biological and Environmental Research under Award Number DE-SC-0023103 and Department of Energy Contract DE-AC36-08GO28308, SUB-2023-10388. M. H. Chou is supported by the Herman F. Heep and Minnie Belle Heep Texas A&M University Endowed Fund, held and administered by the Texas A&M Foundation.

