## Supplementary Information for "Evaluating Limits of Machine Learning-Assisted Raman Spectroscopy in Classification of Biological Samples"

**Figure S22** Row-wise normalized confusion matrix for multiclass classification of inter-day data using Neural Network. 23
**Figure S23** Row-wise normalized confusion matrix for multiclass classification of inter-day data using Convolutional Neural Network. 23
**Figure S24** Mean spectra and double standard deviations (shaded region) for E. coli, EBY100, L. lactis, L. reuteri, YAG01, YAG02, YAG03, YAG04, YAG05, YAG06, YAG07, YAG08, YAG09, YAG10, YAG17, YAG20, YAG22, YAG23, YAG28, and YLH2. 27
**Figure S25** Binary classification results for all the cells compared to YLH2 using Naïve Bayes Gaussian. 28
**Figure S26** Binary classification results for all the cells compared to YLH2 using Decision Tree. 28
**Figure S27** Binary classification results for all the cells compared to YLH2 using KNN-Euclidean. 29
**Figure S28** Binary classification results for all the cells compared to YLH2 using Neural Network. 29
**Figure S29** Binary classification results for all the cells compared to YLH2 using Convolutional Neural Network. 30
**Figure S30** Row-wise normalized confusion matrix for multiclass classification of cells using Naïve Bayes Gaussian (Overall accuracy: 52.08%). 31
**Figure S31** Row-wise normalized confusion matrix for multiclass classification of cells using a Decision Tree (Overall accuracy: 54.79%). 32
**Figure S32** Row-wise normalized confusion matrix for multiclass classification of cells using KNN-Euclidean (Overall accuracy: 55.76%). 33
**Figure S33** Row-wise normalized confusion matrix for multiclass classification of cells using a Neural network (Overall accuracy- 62.29%). 34
**Figure S34** Row-wise normalized confusion matrix for multiclass classification of cells using a Convolutional Neural Network (Overall accuracy- 68.75%). 35

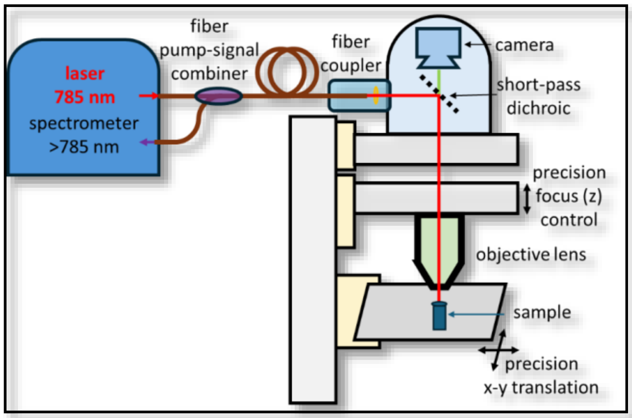

**Figure S1** Custom-assembled Raman microscope.

| GTO (Vol %) | OA (Vol %) |
|---|---|
| 25 | 75 |
| 50 | 50 |
| 75 | 25 |
| 90 | 10 |
| 95 | 5 |
| 97.5 | 2.5 |
| 98.75 | 1.25 |
| 99.375 | 0.625 |
| 99.6875 | 0.3125 |
| 99.8438 | 0.1562 |
| 99.9219 | 0.0781 |
| 99.9609 | 0.0391 |
| 99.9805 | 0.0195 |

**Table S1** Composition (vol%) of mixtures prepared using the weighted average of the spectra from pure Glyceryl Trioctanoate (GTO) and pure Octanoic Acid (OA).

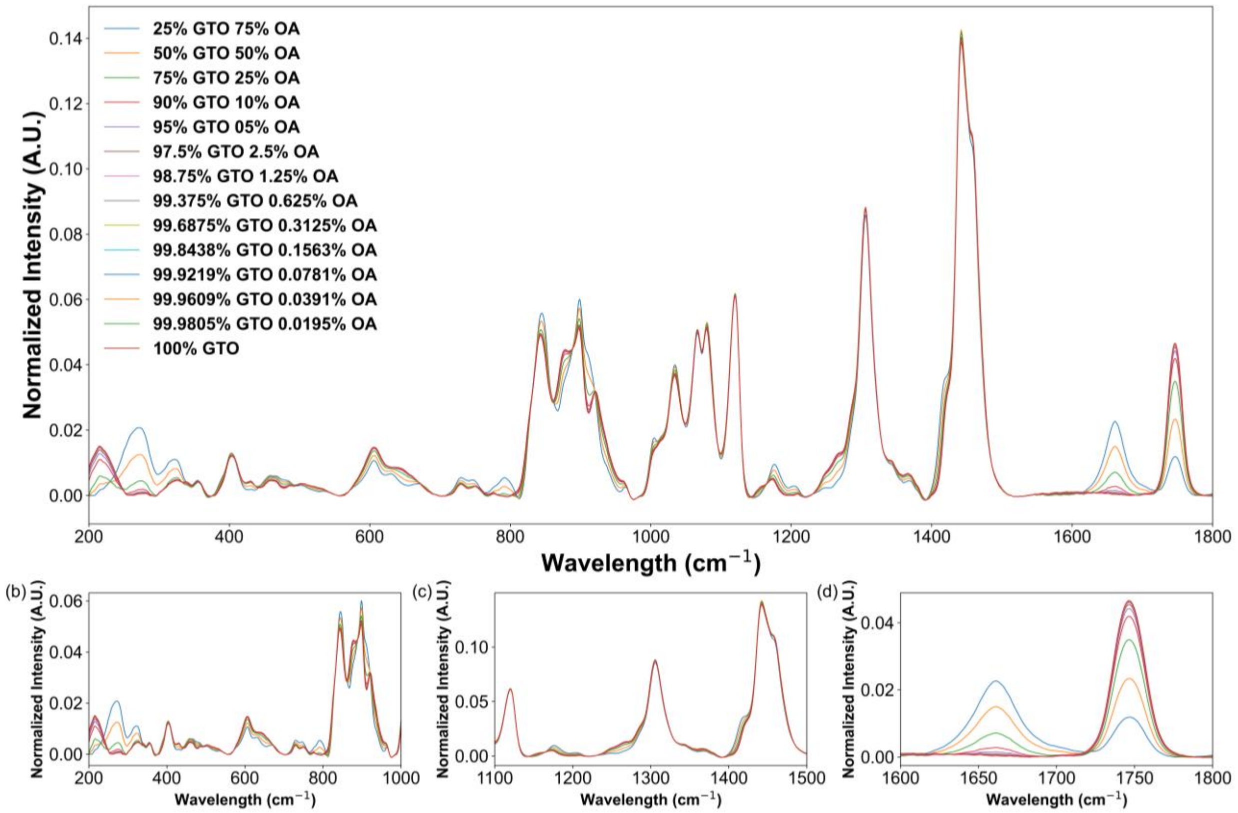

**Figure S2** (a) Normalized Raman spectra for different compositions of Glyceryl trioctanoate (GTO) and Octanoic acid (OA) generated using a weighted average of pure GTO and OA. (b) 200–1000 $cm^{-1}$, (c) 1100–1500 $cm^{-1}$, and (d) 1600–1800 $cm^{-1}$ are enlarged versions of specific regions, highlighting subtle spectral variations with changing OA concentration.

| **Mixture Compositions** | **Similarity Metrics** | |
|---|---|---|
| | **PC** | **PD** |
| **25% GTO 75% OA** | 0.952229 | 0.034831 |
| **50% GTO 50% OA** | 0.97898 | 0.02335 |
| **75% GTO 25% OA** | 0.994218 | 0.011825 |
| **90% GTO 10% OA** | 0.99897 | 0.004919 |
| **95% GTO 5% OA** | 0.999679 | 0.002752 |
| **97.5% GTO 2.5% OA** | 0.999872 | 0.00168 |
| **98.75% GTO 1.25% OA** | 0.999923 | 0.001121 |
| **99.375% GTO 0.625% OA** | 0.999936 | 0.000899 |
| **99.6875% GTO 0.3125% OA** | 0.999939 | 0.00085 |
| **99.8438% GTO 0.1563% OA** | 0.99994 | 0.000839 |
| **99.9219% GTO 0.0781% OA** | 0.99994 | 0.000837 |
| **99.9609% GTO 0.0391% OA** | 0.99994 | 0.000836 |
| **99.9805% GTO 0.0195% OA** | 0.99994 | 0.000836 |

**Table S2** Similarity score for spectra with different binary mixtures of GTO and OA compared to pure GTO (100%) using the two similarity metrics for average standard deviation $1.67\times10^{-4}$ ($\sigma$=0.5). Green indicates high similarity, while red represents dissimilarity.

| Noise Levels (σ) | Average Standard Deviation |
|---|---|
| **0.5** | $1.67\times10^{-4}$ |
| **1** | $3.33\times10^{-4}$ |
| **2** | $6.64\times10^{-4}$ |
| **5** | $1.62\times10^{-3}$ |
| **7** | $2.23\times10^{-3}$ |
| **10** | $3.08\times10^{-3}$ |
| **15** | $4.33\times10^{-3}$ |
| **45** | $8.32\times10^{-3}$ |
| **60** | $9.14\times10^{-3}$ |

**Table S3** Average standard deviation for each noise level for the simulated spectra.

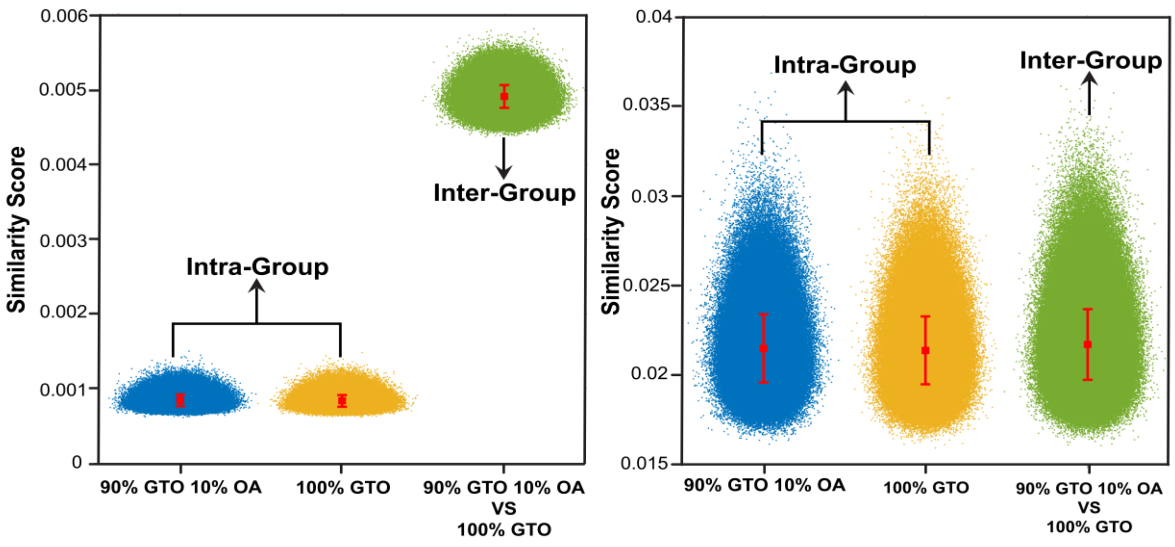

**Figure S3** Scatter plot for similarity score for 90% GTO 10% OA compared to 100% GTO using Pair Distance (PD) for noise levels (σ) (a) 0.5 and (b) 15. As noise increases (σ = 15), intra-group similarity scores become substantially more spread out and overlap with inter-group scores, resulting in reduced classification accuracy. Red dots and error bars represent the mean and standard deviation of similarity scores for each group.

| Mixture Compositions | Average standard deviation | | | | | | | | |
|---|---|---|---|---|---|---|---|---|---|
| | $1.67\times10^{-4}$ (σ=0.5) | $3.33\times10^{-4}$ (σ=1) | $6.64\times10^{-4}$ (σ=2) | $1.62\times10^{-3}$ (σ=5) | $2.23\times10^{-3}$ (σ=7) | $3.08\times10^{-3}$ (σ=10) | $4.33\times10^{-3}$ (σ=15) | $8.32\times10^{-3}$ (σ=45) | $9.14\times10^{-3}$ (σ=60) |
| 25% GTO 75% OA | 100% | 100% | 100% | 100% | 100% | 100% | 100% | 100% | 100% |
| 50% GTO 50% OA | 100% | 100% | 100% | 100% | 100% | 100% | 100% | 100% | 99.2% |
| 75% GTO 25% OA | 100% | 100% | 100% | 100% | 100% | 100% | 100% | 97.65% | 93.95% |
| 90% GTO 10% OA | 100% | 100% | 100% | 100% | 100% | 99.75% | 96.85% | 70% | 59.5% |
| 95% GTO 5% OA | 100% | 100% | 100% | 99.4% | 97.45% | 96.7% | 85.4% | 55.5% | 53.15% |
| 97.5% GTO 2.5% OA | 100% | 100% | 99.95% | 95.7% | 88.15% | 76.65% | 59.5% | 53.55% | 53.4% |
| 98.75% GTO 1.25% OA | 100% | 99.9% | 96.7% | 74.85% | 64.05% | 58.25% | 53.8% | 52.55% | 52.3% |
| 99.375% GTO 0.625% OA | 99.95% | 96.4% | 83.7% | 57.05% | 55.1% | 54.45% | 53.9% | 52.05% | 50.9% |
| 99.6875% GTO 0.3125% OA | 95.6% | 81.95% | 61.45% | 52.1% | 52.05% | 52.4% | 52% | 52.7% | 53.15% |
| 99.8438% GTO 0.1563% OA | 82.25% | 58.55% | 53.95% | 52.4% | 53% | 50.85% | 51.65% | 52.5% | 51.15% |
| 99.9219% GTO 0.0781% OA | 62.75% | 55.05% | 52.15% | 51.5% | 52.9% | 51.85% | 52.65% | 52.3% | 52.55% |
| 99.9609% GTO 0.0391% OA | 53.45% | 53.6% | 50.35% | 51.8% | 50.9% | 51.5% | 52.25% | 51.45% | 52.85% |
| 99.9805% GTO 0.0195% OA | 51.6% | 52.45% | 51.4% | 51.15% | 52.95% | 51.7% | 52.4% | 47.6% | 51.75% |

**Table S4** Classification accuracy scores for different composition differences with varying average standard deviation values using Naive Bayes Gaussian.

| Mixture Compositions | Average standard deviation | | | | | | | | |
|---|---|---|---|---|---|---|---|---|---|
| | $1.67\times10^{-4}$ (σ=0.5) | $3.33\times10^{-4}$ (σ=1) | $6.64\times10^{-4}$ (σ=2) | $1.62\times10^{-3}$ (σ=5) | $2.23\times10^{-3}$ (σ=7) | $3.08\times10^{-3}$ (σ=10) | $4.33\times10^{-3}$ (σ=15) | $8.32\times10^{-3}$ (σ=45) | $9.14\times10^{-3}$ (σ=60) |
| 25% GTO 75% OA | 100% | 100% | 100% | 100% | 100% | 100% | 100% | 100% | 100% |
| 50% GTO 50% OA | 100% | 100% | 100% | 100% | 100% | 100% | 100% | 100% | 99.15% |
| 75% GTO 25% OA | 100% | 100% | 100% | 100% | 100% | 100% | 100% | 84.05% | 78.05% |
| 90% GTO 10% OA | 100% | 100% | 100% | 100% | 100% | 99.7% | 94.45% | 61.3% | 53.95% |
| 95% GTO 5% OA | 100% | 100% | 100% | 99.65% | 95.55% | 86.05% | 74.1% | 52.7% | 52.55% |
| 97.5% GTO 2.5% OA | 100% | 100% | 99.85% | 84.25% | 73.75% | 64.45% | 54.4% | 51.9% | 52.5% |
| 98.75% GTO 1.25% OA | 100% | 99.75% | 92.15% | 61.3% | 55.65% | 53.95% | 52.05% | 52.3% | 51.6% |
| 99.375% GTO 0.625% OA | 99.9% | 94.5% | 68.75% | 54.6% | 52.9% | 52.75% | 53.8% | 52.85% | 52.65% |
| 99.6875% GTO 0.3125% OA | 91.6% | 68.8% | 53.8% | 51.95% | 52.3% | 52.15% | 53.35% | 52.65% | 52.75% |
| 99.8438% GTO 0.1563% OA | 71.2% | 54.55% | 52.25% | 52.1% | 53.65% | 51.4% | 51.9% | 52.15% | 52.2% |
| 99.9219% GTO 0.0781% OA | 55.05% | 51.9% | 51.85% | 51.7% | 51.7% | 52.3% | 51.75% | 52.65% | 52.2% |
| 99.9609% GTO 0.0391% OA | 51.55% | 53.35% | 51.4% | 52.9% | 51.45% | 51.75% | 52.45% | 52.25% | 53.05% |
| 99.9805% GTO 0.0195% OA | 52.65% | 52.15% | 52.6% | 53.05% | 52.1% | 51.75% | 51.55% | 52.45% | 52.85% |

**Table S5** Classification accuracy scores for different composition differences with varying average standard deviation values using a Decision Tree.

| Mixture Compositions | Average standard deviation | | | | | | | | |
|---|---|---|---|---|---|---|---|---|---|
| | $1.67\times10^{-4}$ (σ=0.5) | $3.33\times10^{-4}$ (σ=1) | $6.64\times10^{-4}$ (σ=2) | $1.62\times10^{-3}$ (σ=5) | $2.23\times10^{-3}$ (σ=7) | $3.08\times10^{-3}$ (σ=10) | $4.33\times10^{-3}$ (σ=15) | $8.32\times10^{-3}$ (σ=45) | $9.14\times10^{-3}$ (σ=60) |
| 25% GTO 75% OA | 100% | 100% | 100% | 100% | 100% | 100% | 100% | 100% | 100% |
| 50% GTO 50% OA | 100% | 100% | 100% | 100% | 100% | 100% | 100% | 100% | 100% |
| 75% GTO 25% OA | 100% | 100% | 100% | 100% | 100% | 100% | 100% | 98.6% | 93.85% |
| 90% GTO 10% OA | 100% | 100% | 100% | 100% | 100% | 100% | 99.65% | 71% | 60.4% |
| 95% GTO 5% OA | 100% | 100% | 100% | 99.95% | 99.8% | 97.85% | 86.75% | 56.7% | 53.35% |
| 97.5% GTO 2.5% OA | 100% | 100% | 100% | 97.3% | 88.65% | 77.5% | 59.4% | 52.8% | 52.5% |
| 98.75% GTO 1.25% OA | 100% | 100% | 99.1% | 75.75% | 65.5% | 59.15% | 54.8% | 53.15% | 52.95% |
| 99.375% GTO 0.625% OA | 100% | 98.95% | 83.95% | 56.9% | 54.45% | 55.5% | 54.45% | 52.4% | 50.4% |
| 99.6875% GTO 0.3125%OA | 99% | 82.75% | 62.4% | 53.8% | 51% | 52.2% | 52% | 51.8% | 53.3% |
| 99.8438% GTO 0.1563%OA | 83.2% | 58.5% | 53.7% | 53.25% | 50.1% | 51.5% | 51.05% | 53.1% | 52.55% |
| 99.9219% GTO 0.0781% OA | 62.95% | 55.15% | 52.55% | 52% | 52.65% | 52.4% | 52.85% | 53.35% | 51.75% |
| 99.9609% GTO 0.0391% OA | 53.95% | 54.1% | 49.85% | 53.35% | 51.55% | 52.4% | 51.95% | 50.9% | 53.35% |
| 99.9805% GTO 0.0195% OA | 52% | 52.25% | 50.9% | 50.95% | 52.9% | 52.2% | 52.8% | 47.6% | 52.85% |

**Table S6** Classification accuracy scores for different composition differences with varying average standard deviation values using SVM.

| Mixture Compositions | Average standard deviation | | | | | | | | |
|---|---|---|---|---|---|---|---|---|---|
| | $1.67\times10^{-4}$ (σ=0.5) | $3.33\times10^{-4}$ (σ=1) | $6.64\times10^{-4}$ (σ=2) | $1.62\times10^{-3}$ (σ=5) | $2.23\times10^{-3}$ (σ=7) | $3.08\times10^{-3}$ (σ=10) | $4.33\times10^{-3}$ (σ=15) | $8.32\times10^{-3}$ (σ=45) | $9.14\times10^{-3}$ (σ=60) |
| 25% GTO 75% OA | 100% | 100% | 100% | 100% | 100% | 100% | 100% | 100% | 100% |
| 50% GTO 50% OA | 100% | 100% | 100% | 100% | 100% | 100% | 100% | 100% | 99.65% |
| 75% GTO 25% OA | 100% | 100% | 100% | 100% | 100% | 100% | 100% | 95.1% | 86.3% |
| 90% GTO 10% OA | 100% | 100% | 100% | 100% | 100% | 99.9% | 97.35% | 59.2% | 53.3% |
| 95% GTO 5% OA | 100% | 100% | 100% | 99.9% | 98.15% | 92.9% | 73.15% | 53% | 53.3% |
| 97.5% GTO 2.5% OA | 100% | 100% | 100% | 90.8% | 79.15% | 64.9% | 54.2% | 53.65% | 51.45% |
| 98.75% GTO 1.25% OA | 100% | 99.95% | 95.6% | 62% | 56.05% | 53.75% | 53.15% | 52.7% | 52.65% |
| 99.375% GTO 0.625% OA | 99.95% | 95.3% | 71.15% | 52.55% | 52.95% | 52.45% | 53.35% | 52.6% | 51.05% |
| 99.6875% GTO 0.3125% OA | 94.6% | 68.7% | 55.6% | 52.85% | 51.8% | 52.9% | 52.25% | 52.2% | 51.2% |
| 99.8438% GTO 0.1563% OA | 70.35% | 54.9% | 52.15% | 52.4% | 51.9% | 51.8% | 53.05% | 53.75% | 52.05% |
| 99.9219% GTO 0.0781% OA | 55.05% | 53.5% | 52.4% | 51.25% | 52.65% | 51% | 52% | 52.75% | 52.05% |
| 99.9609% GTO 0.0391% OA | 53.65% | 52.2% | 52.1% | 51% | 52.35% | 51.85% | 51.8% | 54.85% | 51.8% |
| 99.9805% GTO 0.0195% OA | 52.4% | 52.75% | 51.4% | 52.9% | 52.75% | 52.05% | 51.7% | 51.6% | 51.45% |

**Table S7** Classification accuracy scores for different composition differences with varying average standard deviation values using KNN-Euclidean.

| Mixture Compositions | Average standard deviation | | | | | | | | |
|---|---|---|---|---|---|---|---|---|---|
| | $1.67\times10^{-4}$ (σ=0.5) | $3.33\times10^{-4}$ (σ=1) | $6.64\times10^{-4}$ (σ=2) | $1.62\times10^{-3}$ (σ=5) | $2.23\times10^{-3}$ (σ=7) | $3.08\times10^{-3}$ (σ=10) | $4.33\times10^{-3}$ (σ=15) | $8.32\times10^{-3}$ (σ=45) | $9.14\times10^{-3}$ (σ=60) |
| 25% GTO 75% OA | 100% | 100% | 100% | 100% | 100% | 100% | 100% | 100% | 100% |
| 50% GTO 50% OA | 100% | 100% | 100% | 100% | 100% | 100% | 100% | 100% | 100% |
| 75% GTO 25% OA | 100% | 100% | 100% | 100% | 100% | 100% | 100% | 98.65% | 93.95% |
| 90% GTO 10% OA | 100% | 100% | 100% | 100% | 100% | 100% | 99.8% | 70.8% | 60.3% |
| 95% GTO 5% OA | 100% | 100% | 100% | 99.95% | 99.85% | 97.9% | 87.1% | 55.85% | 52.8% |
| 97.5% GTO 2.5% OA | 100% | 100% | 100% | 97.35% | 88.85% | 76.45% | 59% | 54.55% | 52.75% |
| 98.75% GTO 1.25% OA | 100% | 100% | 99% | 75.35% | 64.7% | 58.25% | 54.65% | 52.6% | 52.45% |
| 99.375% GTO 0.625% OA | 100% | 99.05% | 83.35% | 56% | 53.95% | 54.45% | 53.35% | 52.85% | 51.4% |
| 99.6875% GTO 0.3125% OA | 98.9% | 82.5% | 61.4% | 52.6% | 52.15% | 52.2% | 52% | 51.85% | 53.3% |
| 99.8438% GTO 0.1563% OA | 82.95% | 59.75% | 54.05% | 51.55% | 52.65% | 52.8% | 52.25% | 53.25% | 52.6% |
| 99.9219% GTO 0.0781% OA | 63.7% | 54.1% | 51.75% | 51.7% | 52.35% | 53.35% | 54.4% | 52.85% | 52.1% |
| 99.9609% GTO 0.0391% OA | 55.15% | 54.55% | 51.2% | 53% | 52% | 51.75% | 52.45% | 53.1% | 53.2% |
| 99.9805% GTO 0.0195% OA | 52.75% | 52.45% | 51.85% | 52.8% | 53.35% | 52.05% | 53% | 49.6% | 53.3% |

**Table S8** Classification accuracy score for different composition differences with varying average standard deviation values using a Neural Network.

| Mixture Compositions | Average standard deviation | | | | | | | | |
|---|---|---|---|---|---|---|---|---|---|
| | $1.67\times10^{-4}$ (σ=0.5) | $3.33\times10^{-4}$ (σ=1) | $6.64\times10^{-4}$ (σ=2) | $1.62\times10^{-3}$ (σ=5) | $2.23\times10^{-3}$ (σ=7) | $3.08\times10^{-3}$ (σ=10) | $4.33\times10^{-3}$ (σ=15) | $8.32\times10^{-3}$ (σ=45) | $9.14\times10^{-3}$ (σ=60) |
| 25% GTO 75% OA | 100% | 100% | 100% | 100% | 100% | 100% | 100% | 100% | 100% |
| 50% GTO 50% OA | 100% | 100% | 100% | 100% | 100% | 100% | 100% | 100% | 100% |
| 75% GTO 25% OA | 100% | 100% | 100% | 100% | 100% | 100% | 100% | 99.2% | 95.9% |
| 90% GTO 10% OA | 100% | 100% | 100% | 100% | 100% | 100% | 100% | 80.45% | 73.15% |
| 95% GTO 5% OA | 100% | 100% | 100% | 100% | 100% | 98.6% | 92.3% | 67.1% | 58.7% |
| 97.5% GTO 2.5% OA | 100% | 100% | 100% | 98.45% | 93.45% | 84.6% | 73.9% | 56.5% | 54.6% |
| 98.75% GTO 1.25% OA | 100% | 100% | 99.7% | 84.8% | 75.3% | 67.7% | 59.65% | 55.7% | 55.05% |
| 99.375% GTO 0.625% OA | 100% | 99.25% | 89.8% | 65.2% | 62.8% | 58.25% | 56% | 55.3% | 53.85% |
| 99.6875% GTO 0.3125% OA | 98.9% | 89.6% | 72% | 56.15% | 55% | 53.45% | 53.8% | 54% | 54.75% |
| 99.8438% GTO 0.1563% OA | 89.55% | 73% | 58.75% | 52.7% | 54.1% | 53.5% | 53.1% | 54.6% | 53.25% |
| 99.9219% GTO 0.0781% OA | 72.3% | 56.3% | 55.15% | 53.7% | 53.55% | 54.75% | 53.4% | 54.5% | 55.35% |
| 99.9609% GTO 0.0391% OA | 57% | 53.2% | 52.85% | 52.35% | 52.55% | 53.5% | 53.25% | 54.8% | 53.45% |
| 99.9805% GTO 0.0195% OA | 54.4% | 52.55% | 53.2% | 52.55% | 53.9% | 53.5% | 53.5% | 53.1% | 54.05% |

**Table S9** Classification accuracy scores for different composition differences with varying average standard deviation values using a Convolutional Neural Network.

| Mixture Compositions | Labels |
|---|---|
| 25% GTO 75% OA | 25% GTO |
| 50% GTO 50% OA | 50% GTO |
| 75% GTO 25% OA | 75% GTO |
| 90% GTO 10% OA | 90% GTO |
| 95% GTO 5% OA | 95% GTO |
| 97.5% GTO 2.5% OA | 97.5% GTO |
| 98.75% GTO 1.25% OA | 98.75% GTO |
| 99.375% GTO 0.625% OA | 99.375% GTO |
| 99.6875% GTO 0.3125% OA | 99.6875% GTO |
| 99.8438% GTO 0.1563% OA | 99.8438% GTO |
| 99.9219% GTO 0.0781% OA | 99.9219% GTO |
| 99.9609% GTO 0.0391% OA | 99.9609% GTO |
| 99.9805% GTO 0.0195% OA | 99.9805% GTO |
| 100% GTO | 100% GTO |

**Table S10** Label details for multiclass classification confusion matrices.

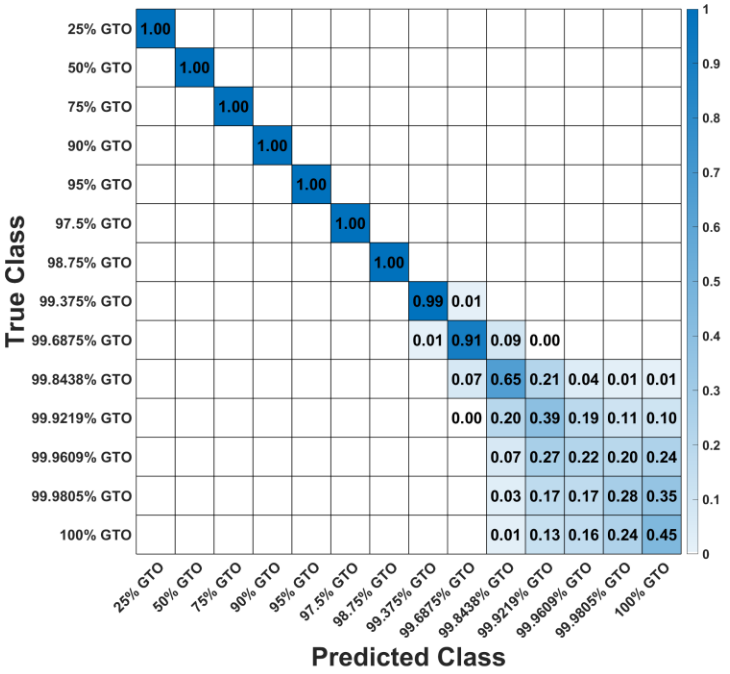

**Figure S4** Row-wise normalized confusion matrix for multiclass classification of simulated mixtures, average standard deviation 1.67×10-4 (σ=0.5), using SVM-Linear.

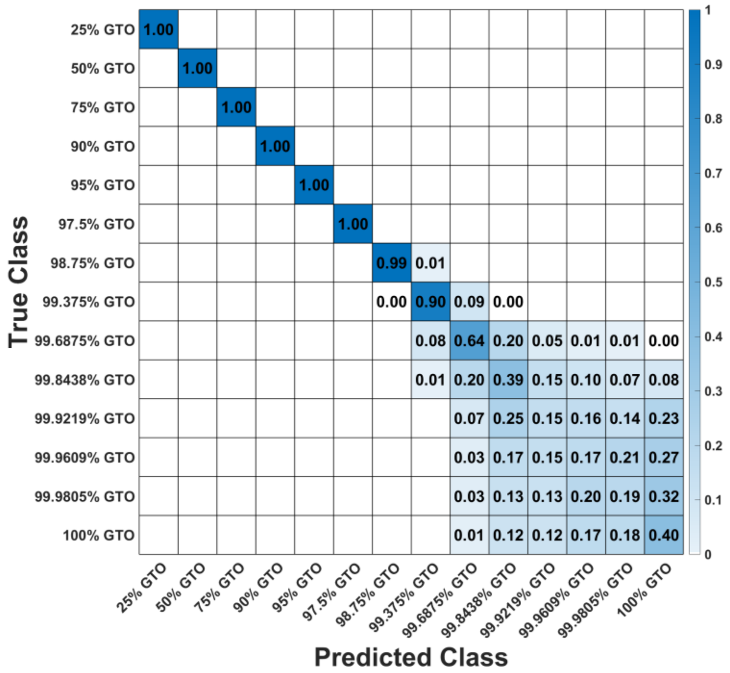

**Figure S5** Row-wise normalized confusion matrix for multiclass classification of simulated mixtures, average standard deviation 3.33×10-4 (σ=1), using SVM-Linear.

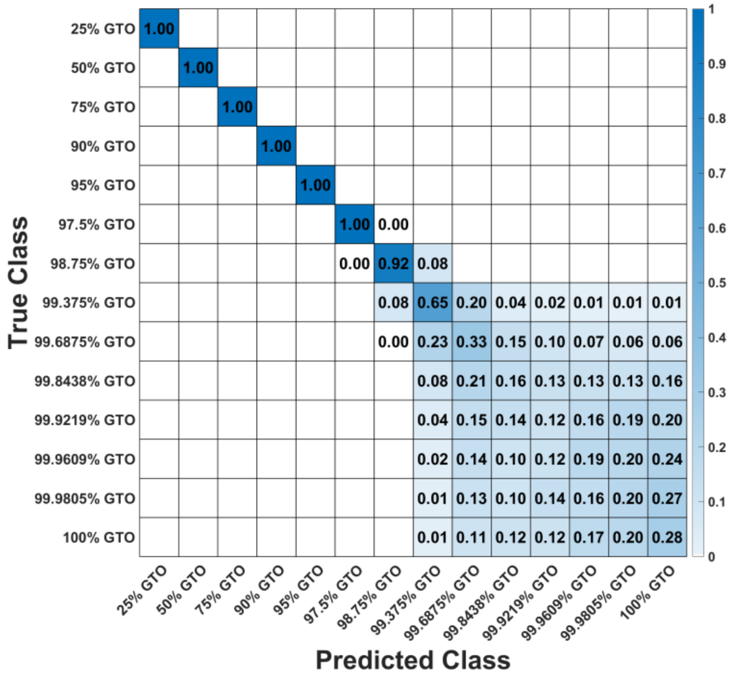

**Figure S6** Row-wise normalized confusion matrix for multiclass classification of simulated mixtures, average standard deviation 6.64×10-4 (σ=2), using SVM-Linear.

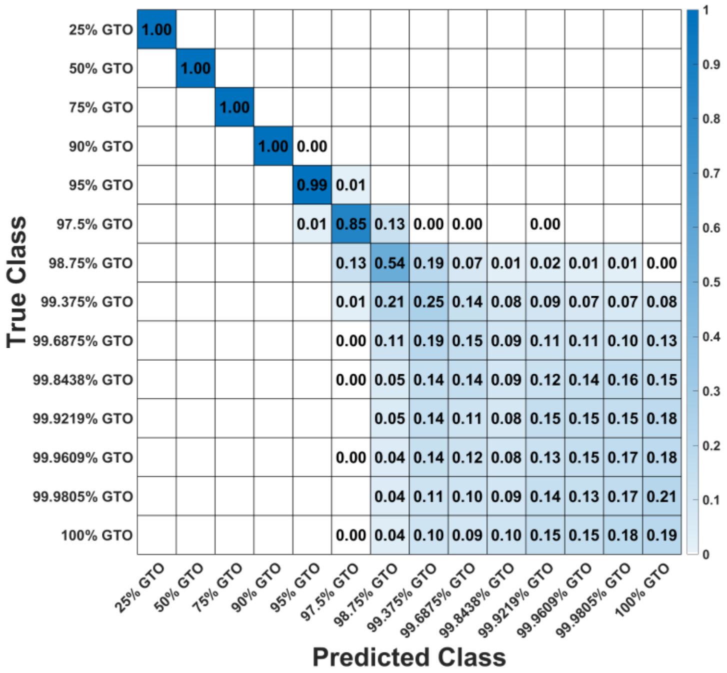

**Figure S7** Row-wise normalized confusion matrix for multiclass classification of simulated mixtures, average standard deviation 1.62×10-3 (σ=5), using SVM-Linear.

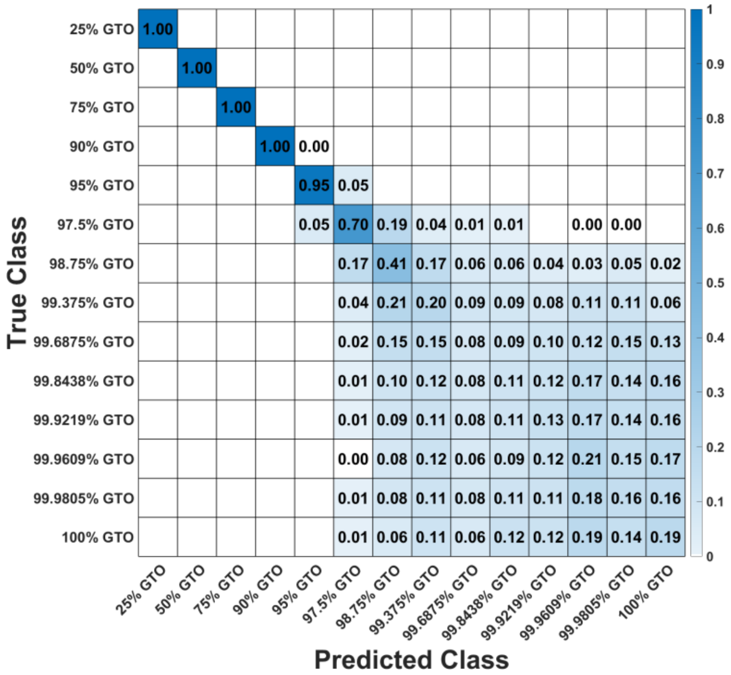

**Figure S8** Row-wise normalized confusion matrix for multiclass classification of simulated mixtures, average standard deviation 2.23×10-3 (σ=7), using SVM-Linear.

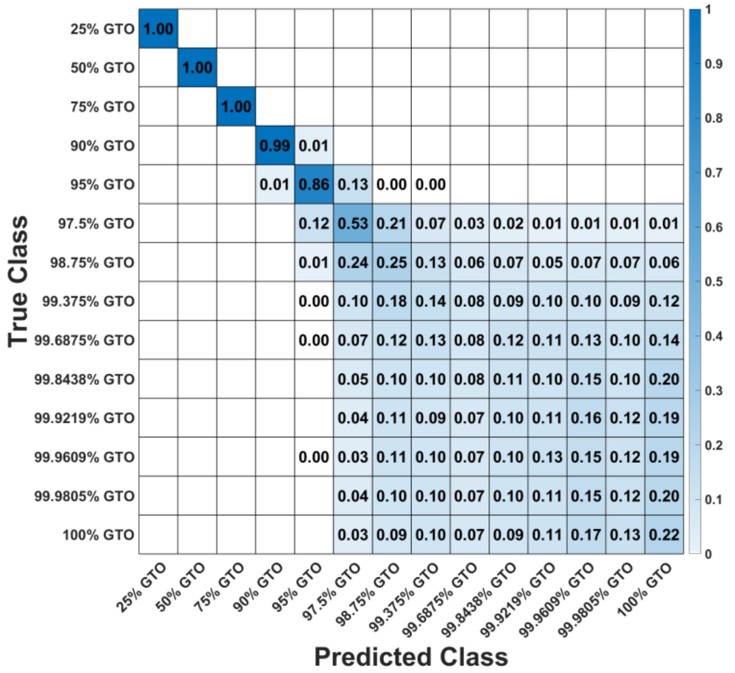

**Figure S9** Row-wise normalized confusion matrix for multiclass classification of simulated mixtures, average standard deviation 3.08×10-3 (σ=10), using SVM-Linear.

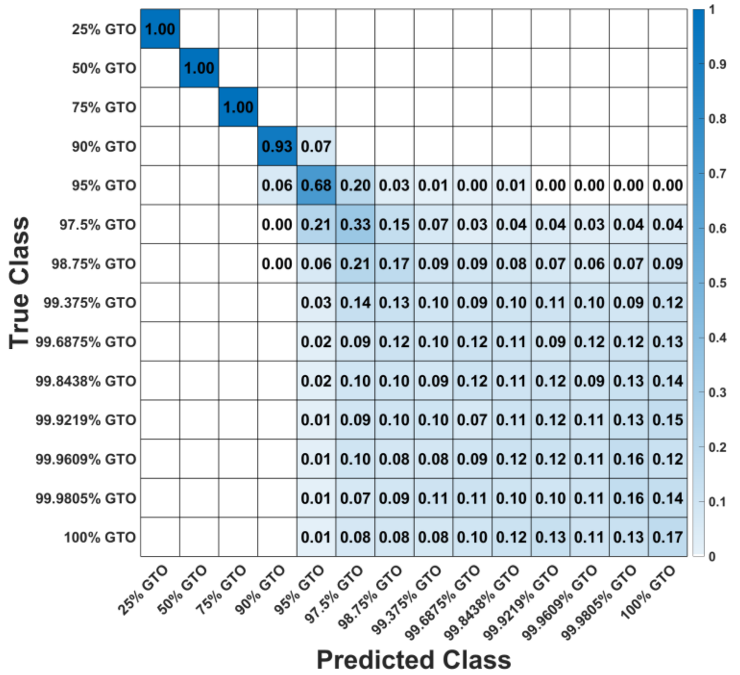

**Figure S10** Row-wise normalized confusion matrix for multiclass classification of simulated mixtures, average standard deviation 4.33×10-3 (σ=15), using SVM-Linear.

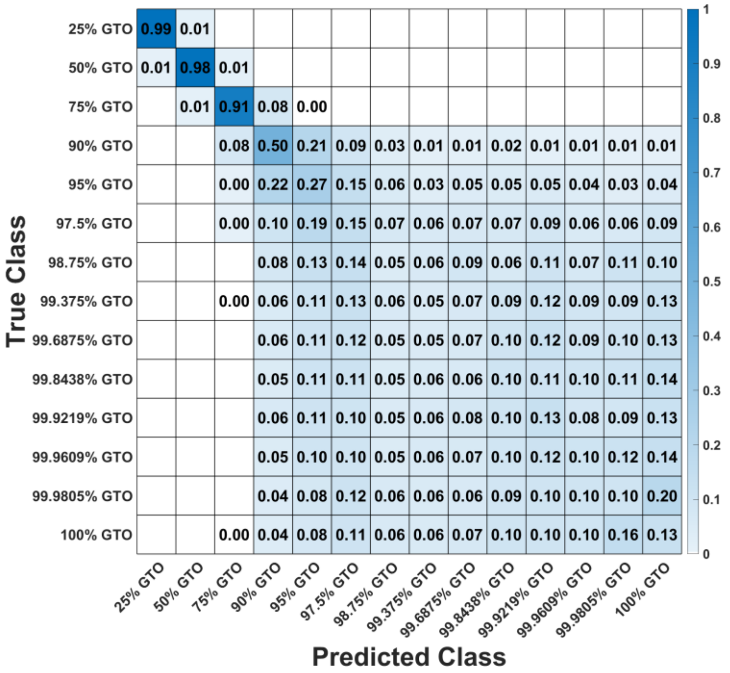

**Figure S11** Row-wise normalized confusion matrix for multiclass classification of simulated mixtures, average standard deviation 8.32×10-3 (σ=45), using SVM-Linear.

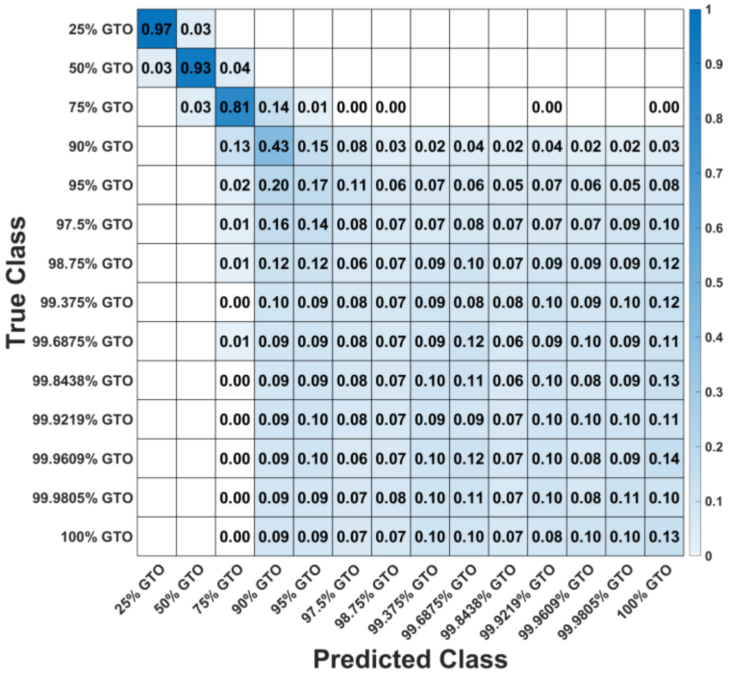

**Figure S12** Row-wise normalized confusion matrix for multiclass classification of simulated mixtures, average standard deviation 9.14×10-3 (σ=60), using SVM-Linear.

| Composition (Vol%) | AVERAGE STANDARD DEVIATION | |
|---|---|---|
| | Inter-day Samples | Intra-day Samples |
| 90% GTO 10 % OA | $4.71\times10^{-4}$ | $4.1\times10^{-4}$ |
| 95% GTO 05 % OA | $4.64\times10^{-4}$ | $4.07\times10^{-4}$ |
| 97.5% GTO 2.5 % OA | $4.82\times10^{-4}$ | $4.08\times10^{-4}$ |
| 98.75% GTO 1.25 % OA | $4.8\times10^{-4}$ | $4.16\times10^{-4}$ |
| 99.375% GTO 0.625 % OA | $4.77\times10^{-4}$ | $4.06\times10^{-4}$ |
| 99.6875% GTO 0.3125 % OA | $4.73\times10^{-4}$ | $4.08\times10^{-4}$ |
| 99.8438% GTO 0.1563 % OA | $4.72\times10^{-4}$ | $4.08\times10^{-4}$ |
| 99.9219% GTO 0.0781 % OA | $4.7\times10^{-4}$ | $4.13\times10^{-4}$ |
| 99.9609% GTO 0.0391 % OA | $4.66\times10^{-4}$ | $4.12\times10^{-4}$ |
| 99.9805% GTO 0.0195 % OA | $4.61\times10^{-4}$ | $4.12\times10^{-4}$ |
| 100% GTO | $4.49\times10^{-4}$ | $4.16\times10^{-4}$ |

**Table S11** Average standard deviation value for inter-day and intra-samples for all the mixtures.

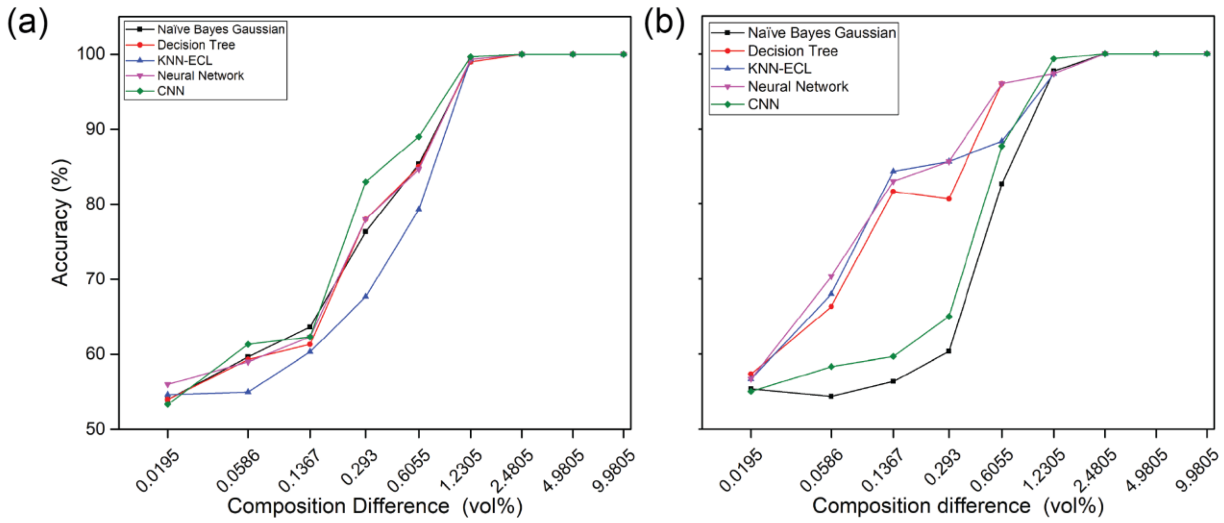

**Figure S13** Classification accuracy for different composition differences (a) Results for intra-day measurements and (b) Results for inter-day measurements.

| Composition difference (vol%) | Accuracy for Simulated Data | Accuracy for Experimental Data |
|---|---|---|
| **9.98046875** | 100% | 100% |
| **4.98046875** | 100% | 100% |
| **2.48046875** | 100% | 100% |
| **1.23046875** | 100% | 99.33% |
| **0.60546875** | 96.1% | 84.66% |
| **0.29296875** | 74.4% | 76.66% |
| **0.13671875** | 56.25% | 66% |
| **0.05859375** | 52.5% | 58.33% |
| **0.01953125** | 52.7% | 52.66% |

**Table S12** Comparison of classification accuracy using a Linear Support Vector Machine (SVM-Linear) for simulated versus experimental intra-day sample data.

| Composition difference (vol%) | Accuracy for Simulated Data | Accuracy for Experimental Data |
|---|---|---|
| **9.98046875** | 100% | 100% |
| **4.98046875** | 100% | 100% |
| **2.48046875** | 100% | 100% |
| **1.23046875** | 100% | 97.66% |
| **0.60546875** | 94.2% | 80.66% |
| **0.29296875** | 69.85% | 63% |
| **0.13671875** | 57.15% | 59% |
| **0.05859375** | 53.05% | 54% |
| **0.01953125** | 53.25% | 57% |

**Table S13** Comparison of classification accuracy using a Linear Support Vector Machine (SVM-Linear) for simulated versus experimental inter-day sample data.

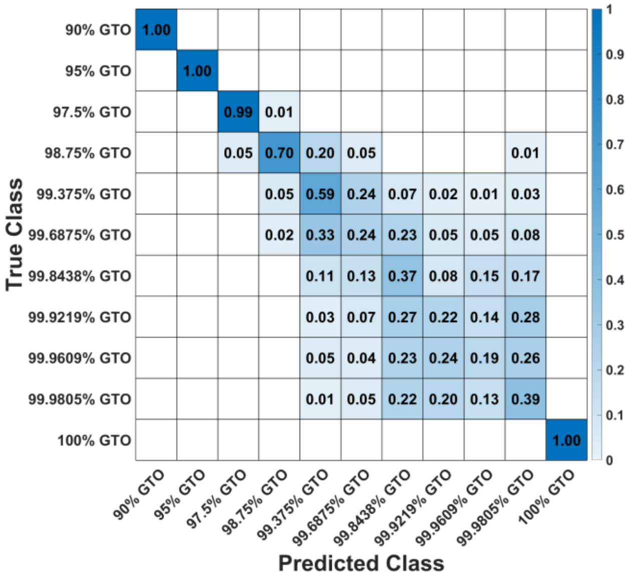

**Figure S14** Row-wise normalized confusion matrix for multiclass classification of intra-day data using Naïve Bayes Gaussian.

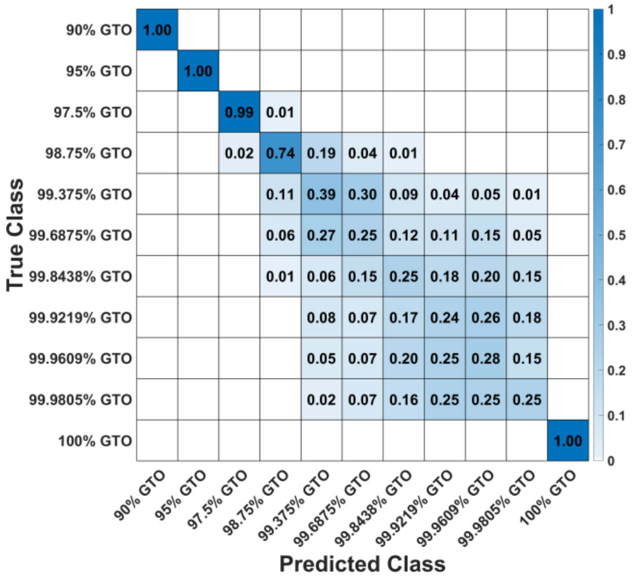

**Figure S15** Row-wise normalized confusion matrix for multiclass classification of intra-day data using Decision Tree.

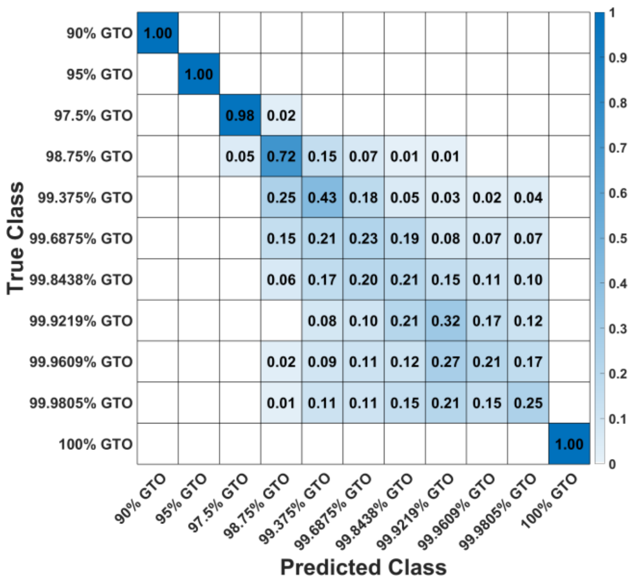

**Figure S16** Row-wise normalized confusion matrix for multiclass classification of intra-day data using KNN-Euclidean.

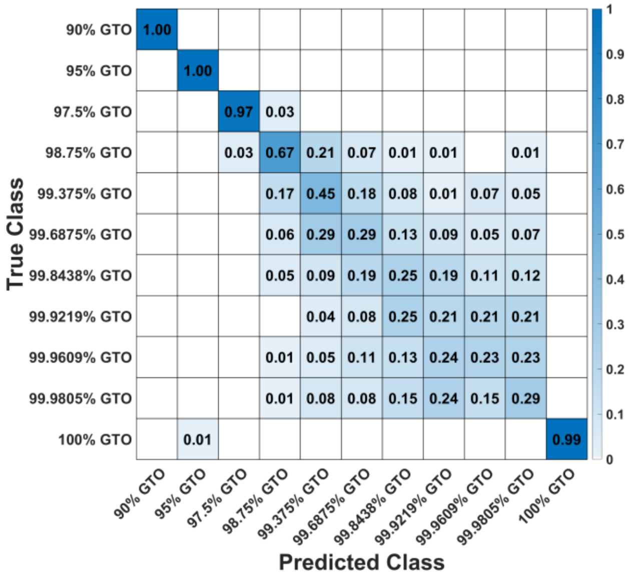

**Figure S17** Row-wise normalized confusion matrix for multiclass classification of intra-day data using Neural Network.

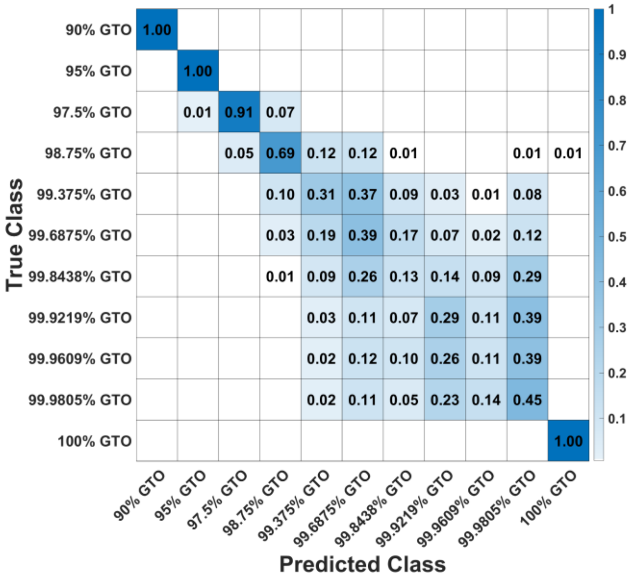

**Figure S18** Row-wise normalized confusion matrix for multiclass classification of intra-day data using Convolutional Neural Network.

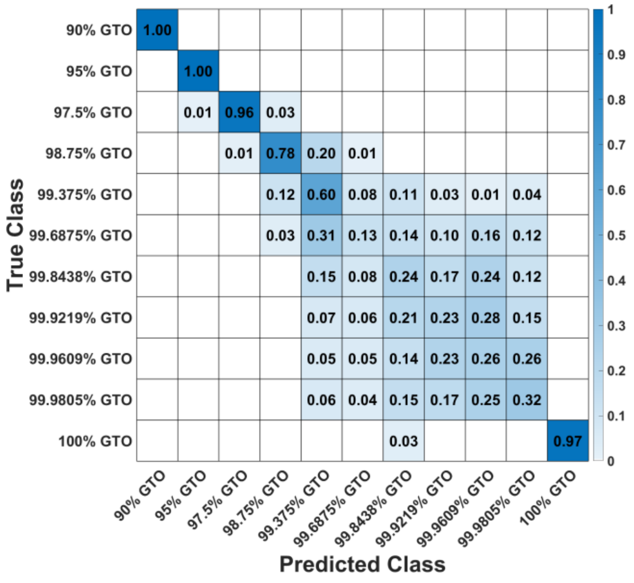

**Figure S19** Row-wise normalized confusion matrix for multiclass classification of inter-day data using Naïve Bayes Gaussian.

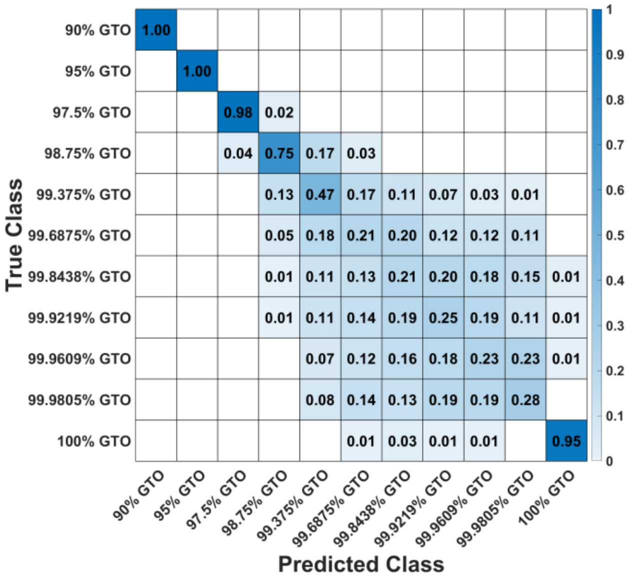

**Figure S20** Row-wise normalized confusion matrix for multiclass classification of inter-day data using Decision Tree.

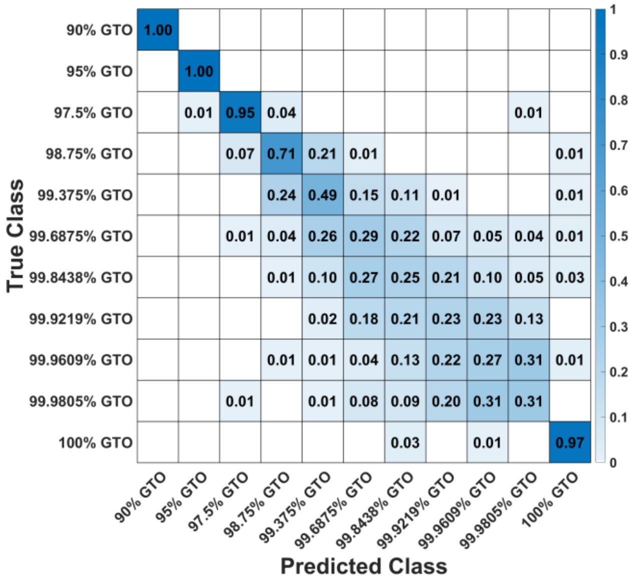

**Figure S21** Row-wise normalized confusion matrix for multiclass classification of inter-day data using KNN-Euclidean.

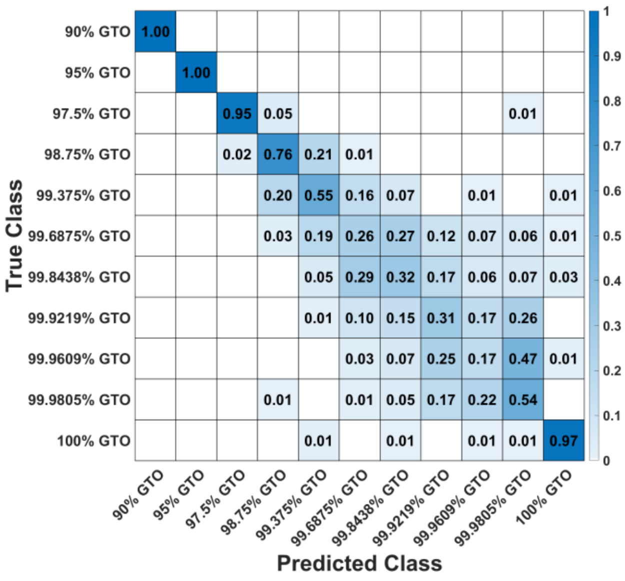

**Figure S22** Row-wise normalized confusion matrix for multiclass classification of inter-day data using Neural Network.

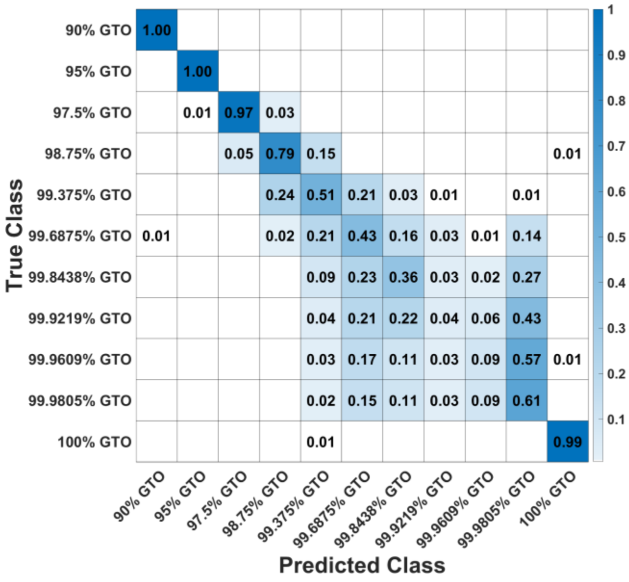

**Figure S23** Row-wise normalized confusion matrix for multiclass classification of inter-day data using Convolutional Neural Network.

| LABELS | GENOTYPE INFORMATION | SOURCE |
| --- | --- | --- |
| *E. coli* | *E. coli* BL21(DE3) $F^-$ *dcm ompT hsdS*($r_B^-$ $m_B^-$) *gal* λ(DE3) | This Paper |
| EBY100 | MATa *AGA1*::*GAL1-AGA1*::*URA3 ura3-52 trp1 le*u2*Δ1 his3Δ200 pep4*::*HIS3 prb1Δ1.6R can1* GAL | Tsai et al. 2009[1] |
| *L. lactis* | MG1363 | This Paper |
| *L. reuteri* | ATCC BAA-2837 (IDCC 3701) | ATCC |
| YLH2 | GSY1136 YIplac211YB/I/E* Δctt1 | Godara et al. 2019 [2] |
| YAG01 | GSY1136 YIplac211YB/I/E* Δctt1, HIS7 389 | Godara et al. 2019 [2] |
| YAG02 | GSY1136 YIplac211YB/I/E* Δctt1, SRO9/GFD2 int | Godara et al. 2019 [2] |
| YAG03 | GSY1136 YIplac211YB/I/E* Δctt1, TYE7 86 | Godara et al. 2019 [2] |
| YAG04 | GSY1136 YIplac211YB/I/E* Δctt1, FLO1 925 | Godara et al. 2019 [2] |
| YAG05 | GSY1136 YIplac211YB/I/E* Δctt1, DAK2/AQY3 int | Godara et al. 2019 [2] |
| YAG06 | GSY1136 YIplac211YB/I/E* Δctt1, SCY1 1836 | Godara et al. 2019 [2] |
| YAG07 | GSY1136 YIplac211YB/I/E* Δctt1, EPL1 1754 | Godara et al. 2019 [2] |
| YAG08 | GSY1136 YIplac211YB/I/E* Δctt1, ALG6 1411 | Godara et al. 2019 [2] |
| YAG09 | GSY1136 YIplac211YB/I/E* Δctt1, MDS3 ins | Godara et al. 2019 [2] |
| YAG10 | GSY1136 YIplac211YB/I/E* Δctt1, YMRCTy1-3 1078 | Godara et al. 2019 [2] |
| YAG17 | GSY1136 YIplac211YB/I/E* Δctt1, ALG6 1411, EPL1 1754 | Godara et al. 2019 [2] |
| YAG20 | GSY1136 YIplac211YB/I/E* Δctt1, EPL1 1754, DAK2/AQY3 int | Godara et al. 2019 [2] |
| YAG22 | GSY1136 YIplac211YB/I/E* Δctt1, ALG6 1411, DAK2/AQY3 int | Godara et al. 2019 [2] |
| YAG23 | GSY1136 YIplac211YB/I/E* Δctt1, ALG6 1411, DAK2/AQY3 int, EPL1 1754 | Godara et al. 2019 [2] |
| YAG28 | GSY1136 YIplac211YB/I/E* Δctt1, ALG6 1411, DAK2/AQY3 int, YMRCTy1-3 1078 | Godara et al. 2019 [2] |

**Table S14** Strain and mutation label details of microorganisms used in the study.

| Label | Average standard deviation |
|---|---|
| E. coli | $1.01 \times 10^{-2}$ |
| EBY100 | $8.38 \times 10^{-3}$ |
| L. lactis | $1.11 \times 10^{-2}$ |
| L. reuteri | $9.37 \times 10^{-3}$ |
| YAG01 | $5.98 \times 10^{-3}$ |
| YAG02 | $4.05 \times 10^{-3}$ |
| YAG03 | $4.32 \times 10^{-3}$ |
| YAG04 | $4.9 \times 10^{-3}$ |
| YAG05 | $4.5 \times 10^{-3}$ |
| YAG06 | $7.02 \times 10^{-3}$ |
| YAG07 | $4.67 \times 10^{-3}$ |
| YAG08 | $5.78 \times 10^{-3}$ |
| YAG09 | $5.25 \times 10^{-3}$ |
| YAG10 | $5.8 \times 10^{-3}$ |
| YAG17 | $4.04 \times 10^{-3}$ |
| YAG20 | $4.48 \times 10^{-3}$ |
| YAG22 | $3.46 \times 10^{-3}$ |
| YAG23 | $4.08 \times 10^{-3}$ |
| YAG28 | $6.84 \times 10^{-3}$ |
| YLH2 | $3.65 \times 10^{-3}$ |

**Table S15** Average standard deviation of Raman spectra in each microorganism.

| Label | PC | PD |
| --- | --- | --- |
| E. coli | -0.09431 | 0.207603 |
| EBY100 | -0.073745 | 0.19612 |
| L. lactis | -0.030124 | 0.202841 |
| L. reuteri | -0.125977 | 0.207085 |
| YAG01 | 0.921848 | 0.037648 |
| YAG02 | 0.95351 | 0.031026 |
| YAG03 | 0.950231 | 0.032904 |
| YAG04 | 0.941429 | 0.034914 |
| YAG05 | 0.952713 | 0.032311 |
| YAG06 | 0.900465 | 0.041817 |
| YAG07 | 0.949683 | 0.031798 |
| YAG08 | 0.93822 | 0.034172 |
| YAG09 | 0.940327 | 0.032547 |
| YAG10 | 0.928058 | 0.040699 |
| YAG17 | 0.953059 | 0.031008 |
| YAG20 | 0.934264 | 0.05119 |
| YAG22 | 0.959237 | 0.0342 |
| YAG23 | 0.955394 | 0.029217 |
| YAG28 | 0.814117 | 0.103241 |

**Table S16** Similarity scores for spectra of various mutated yeast strains and other microorganisms compared to the industrial strain YLH2. Green indicates high similarity, while red represents dissimilarity.

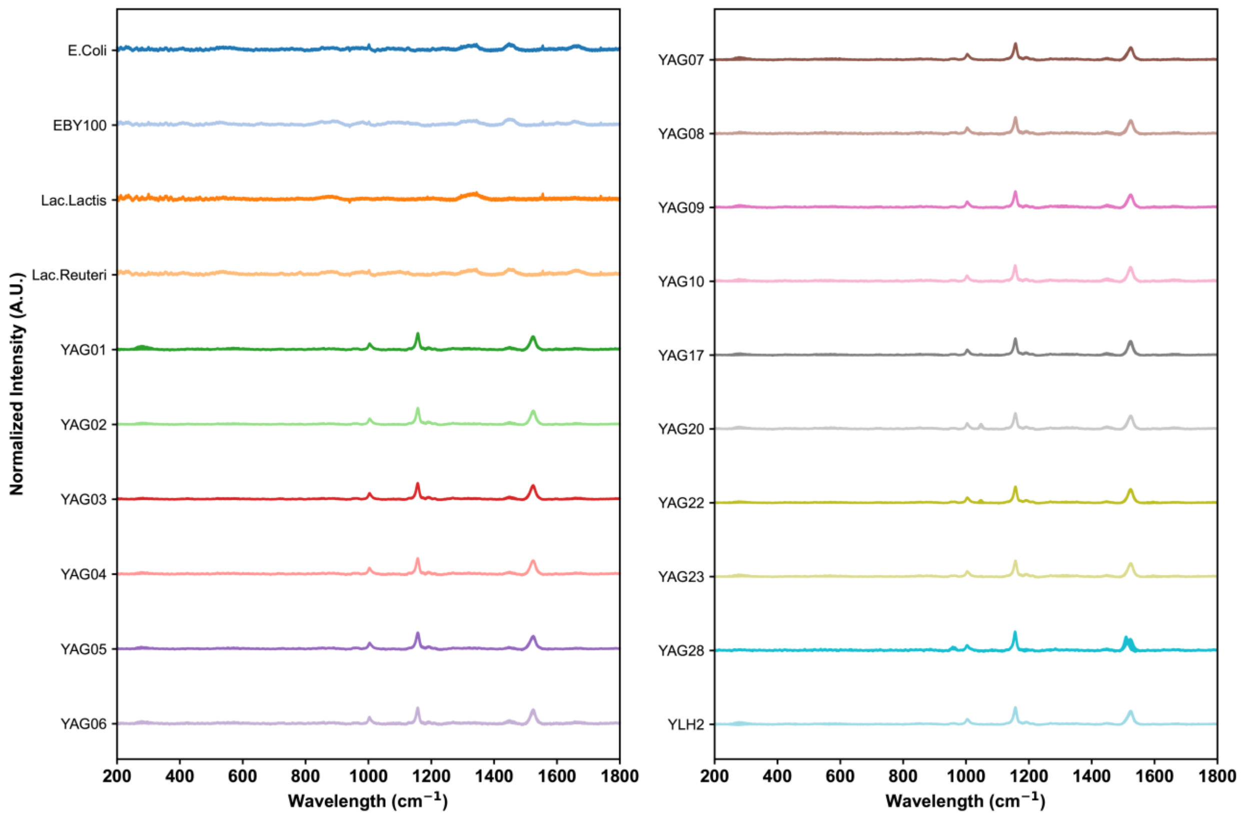

**Figure S24** Mean spectra and double standard deviations (shaded region) for E. coli, EBY100, L. lactis, L. reuteri, YAG01, YAG02, YAG03, YAG04, YAG05, YAG06, YAG07, YAG08, YAG09, YAG10, YAG17, YAG20, YAG22, YAG23, YAG28, and YLH2.

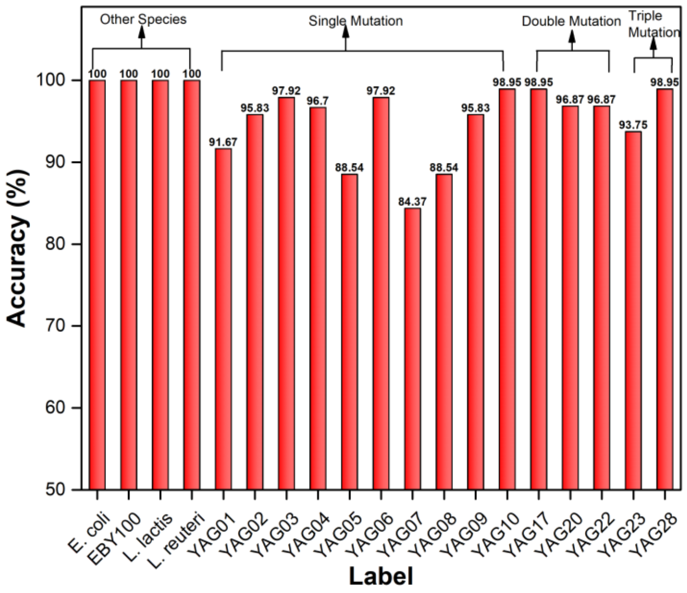

**Figure S25** Binary classification results for all the cells compared to YLH2 using Naïve Bayes Gaussian.

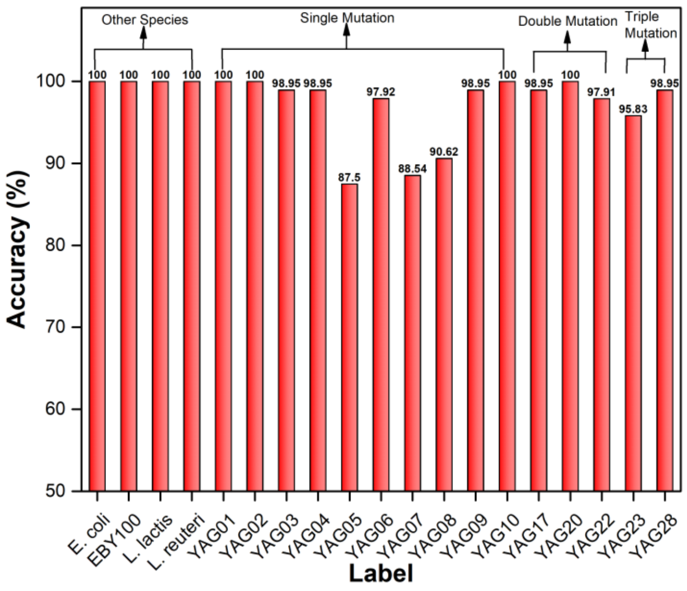

**Figure S26** Binary classification results for all the cells compared to YLH2 using Decision Tree.

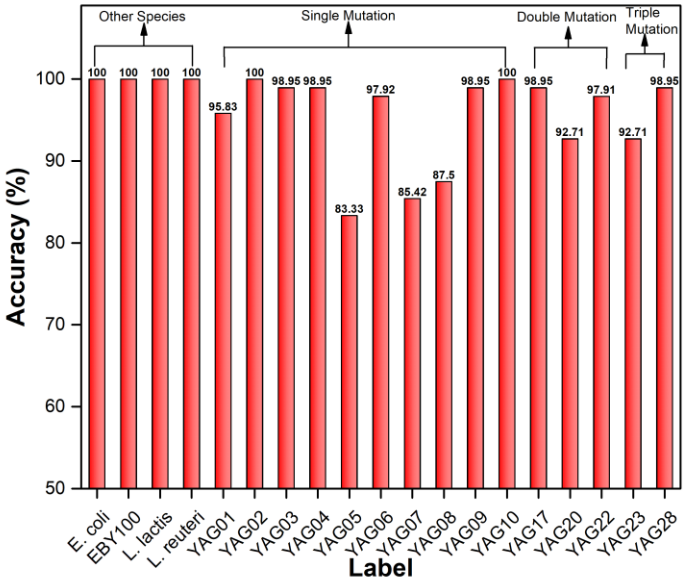

**Figure S27** Binary classification results for all the cells compared to YLH2 using KNN-Euclidean.

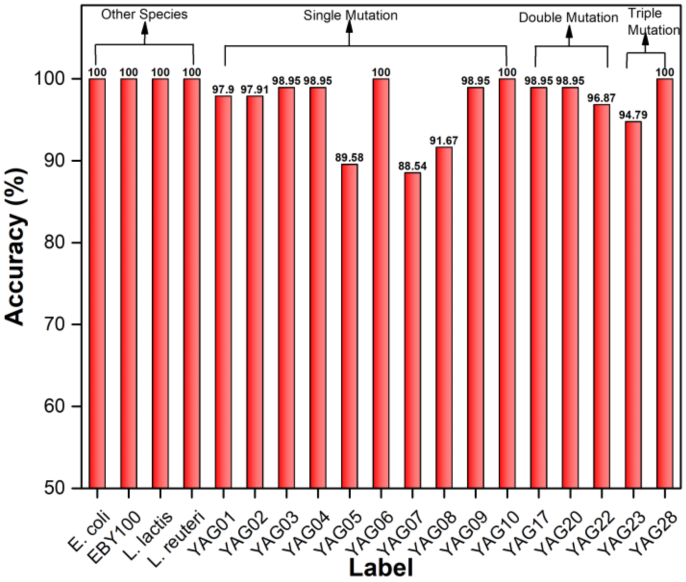

**Figure S28** Binary classification results for all the cells compared to YLH2 using Neural Network.

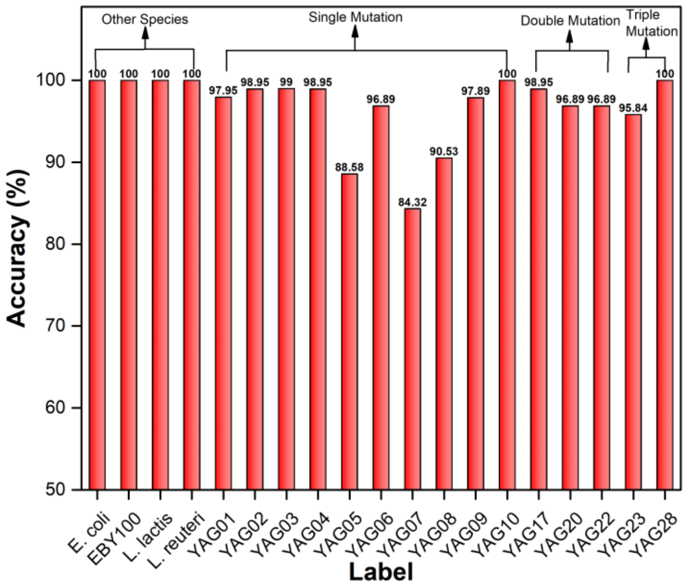

**Figure S29** Binary classification results for all the cells compared to YLH2 using Convolutional Neural Network.

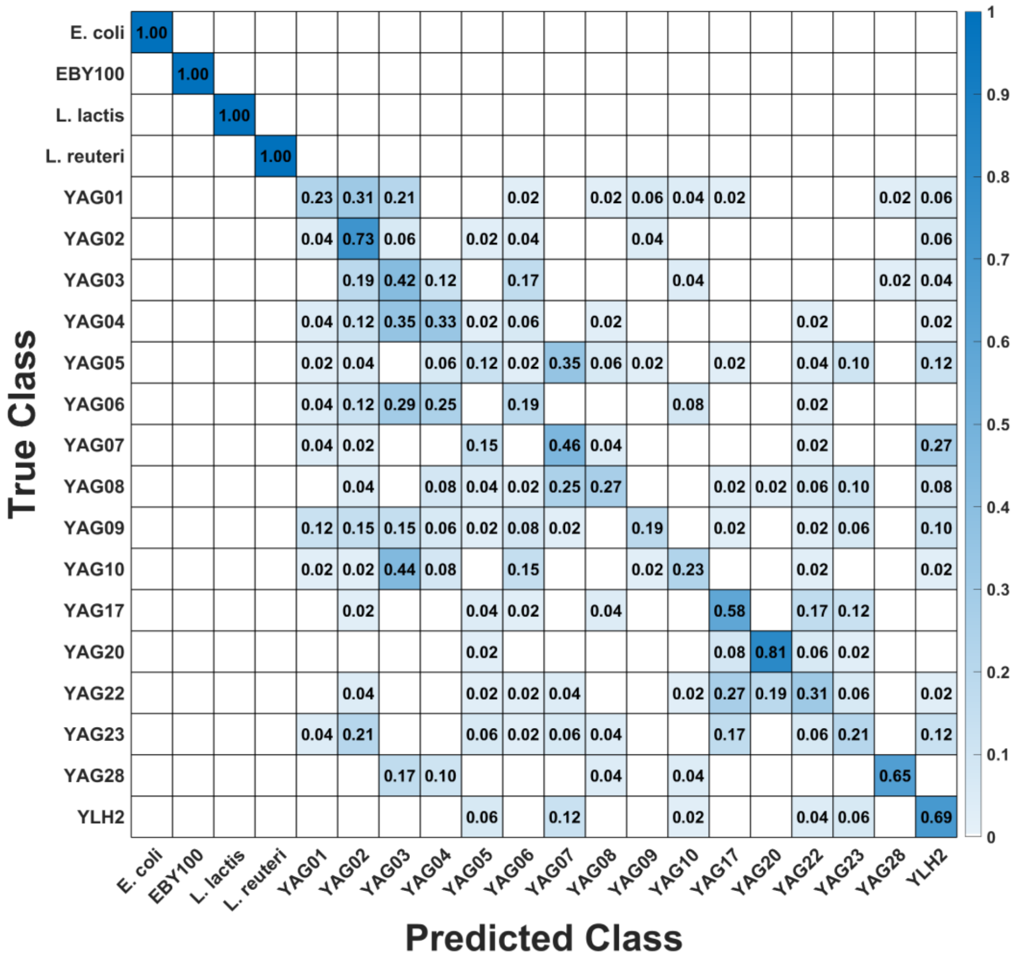

**Figure S30** Row-wise normalized confusion matrix for multiclass classification of cells using Naïve Bayes Gaussian (Overall accuracy: 52.08%).

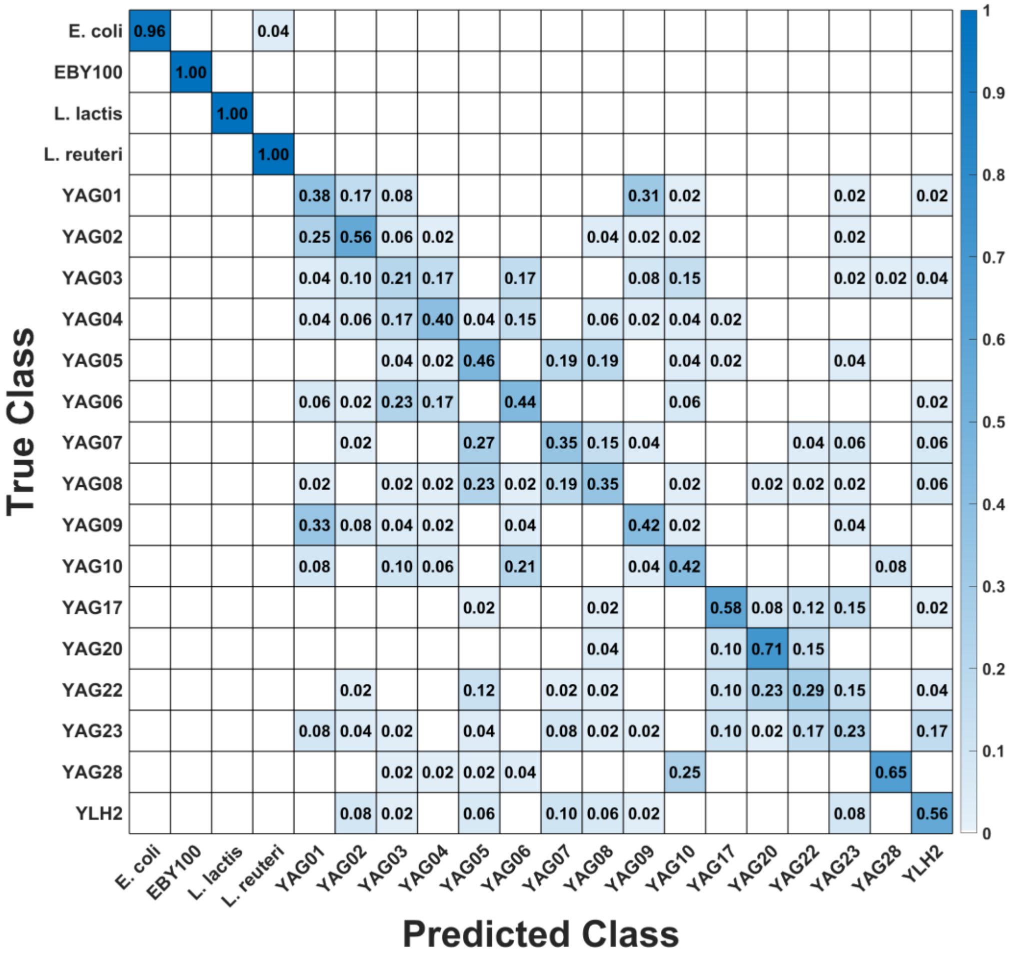

**Figure S31** Row-wise normalized confusion matrix for multiclass classification of cells using a Decision Tree (Overall accuracy: 54.79%).

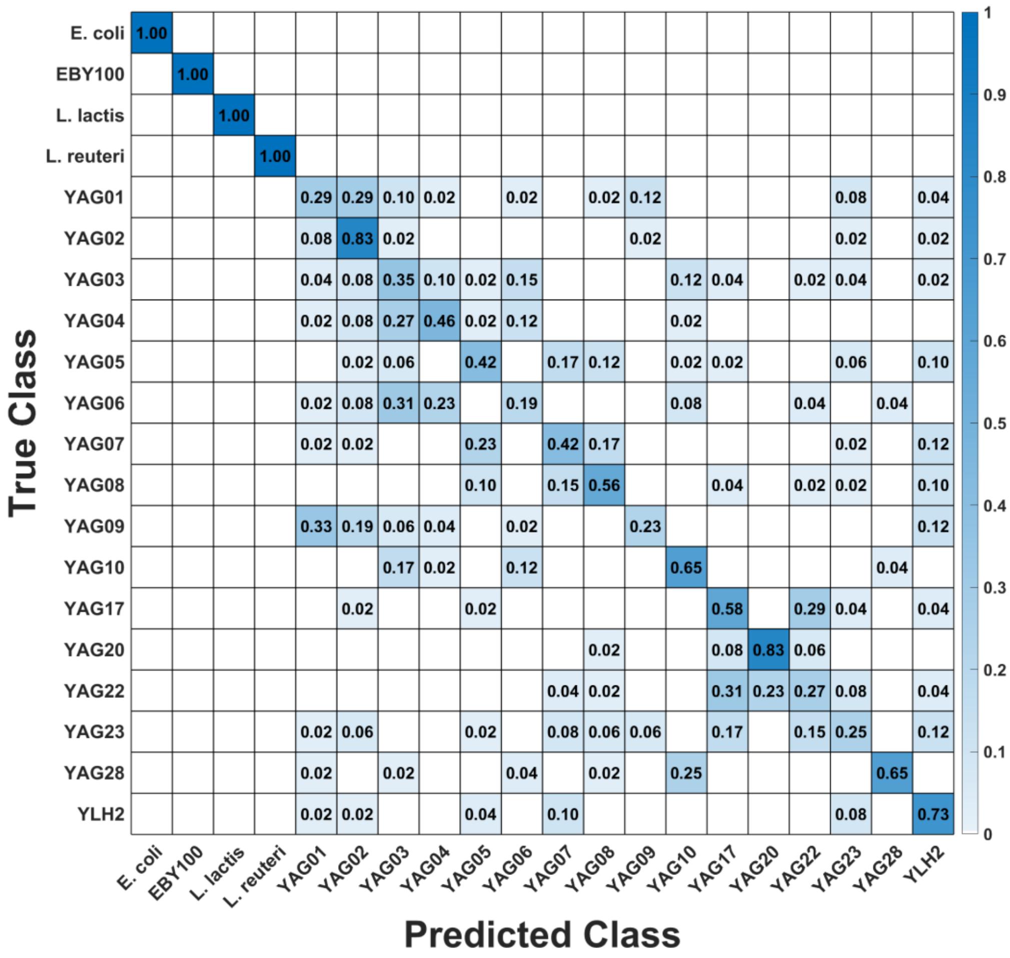

**Figure S32** Row-wise normalized confusion matrix for multiclass classification of cells using KNN-Euclidean (Overall accuracy: 55.76%).

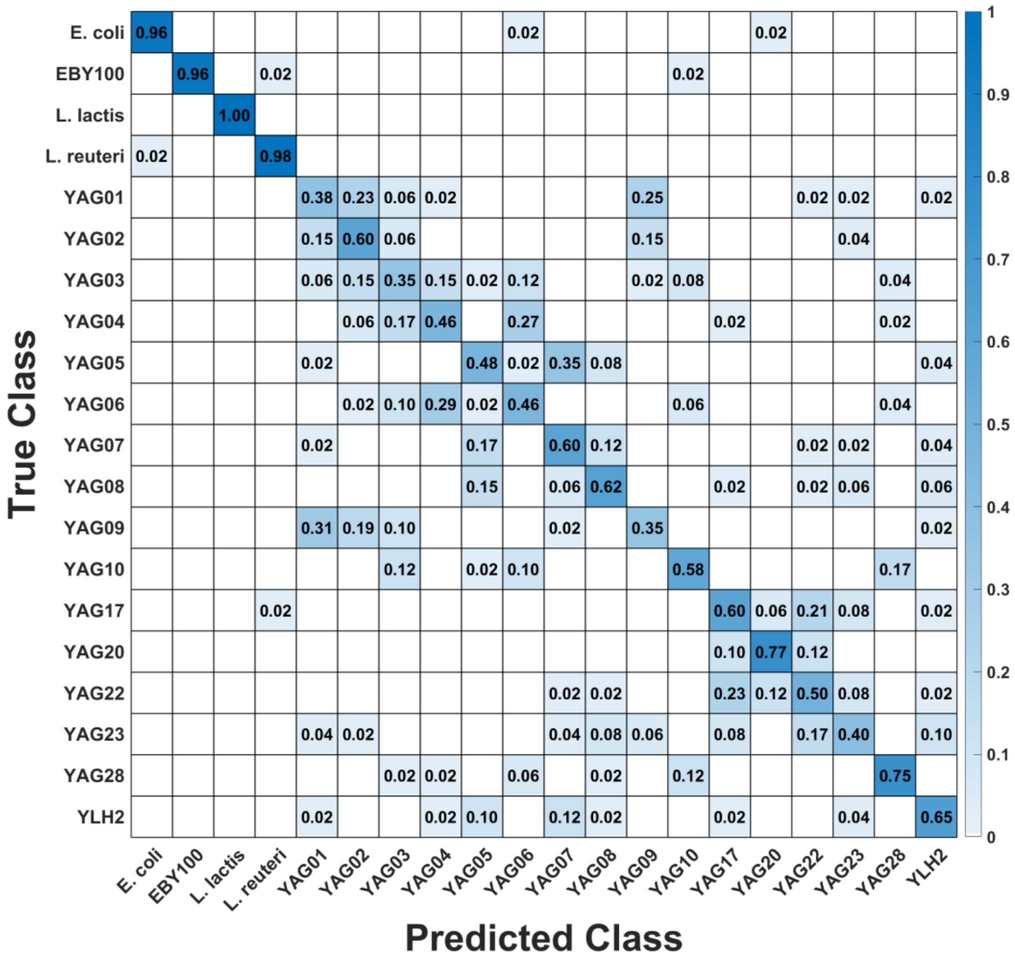

**Figure S33** Row-wise normalized confusion matrix for multiclass classification of cells using a Neural network (Overall accuracy- 62.29%).

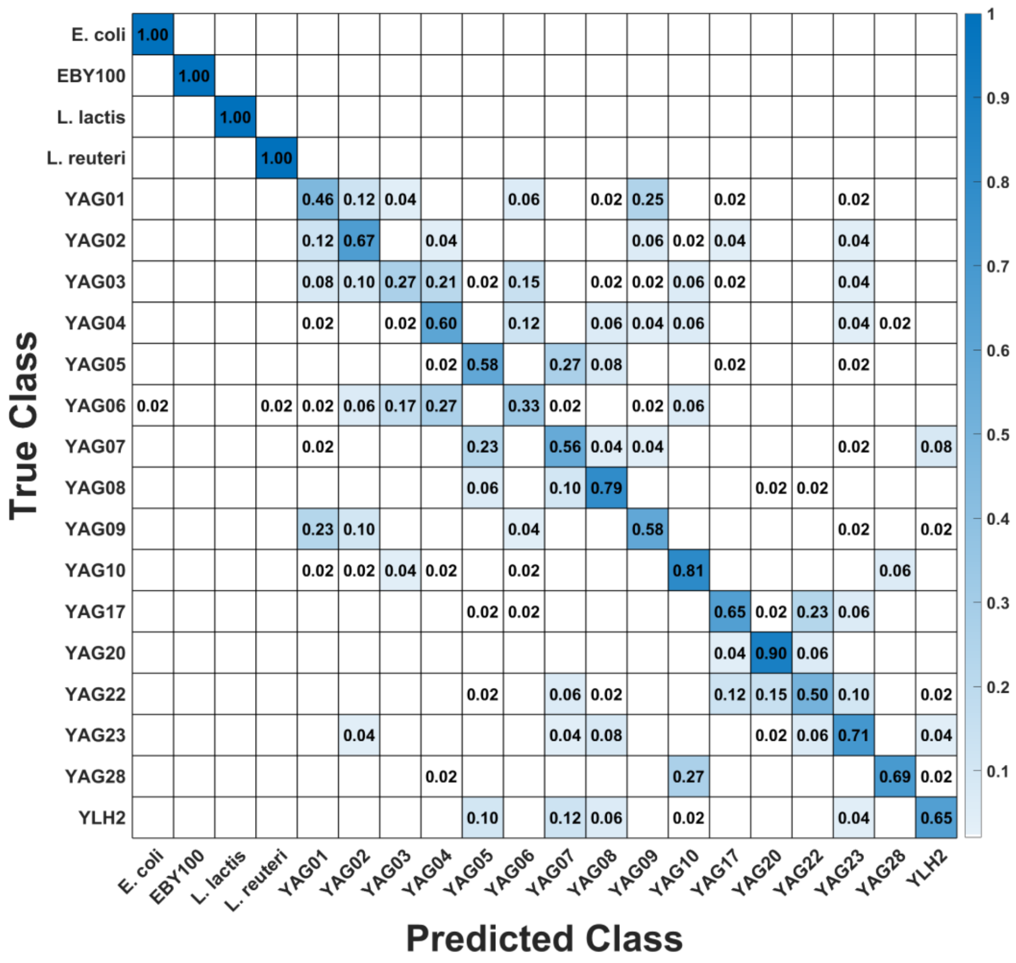

**Figure S34** Row-wise normalized confusion matrix for multiclass classification of cells using a Convolutional Neural Network (Overall accuracy- 68.75%).
